# Changes in DNA Methylation During Anoxia and Reoxygenation in Crucian Carp Brain

**DOI:** 10.1101/2025.07.30.667361

**Authors:** Magdalena Winklhofer, Øivind Andersen, Sjannie Lefevre

**Affiliations:** Section for Physiology and Cell Biology, Department of Biosciences, University of Oslo, Oslo, Norway; Faculty of Biosciences, Department of Animal and Aquacultural Sciences, Genome Biology, Norwegian University of Life Sciences, Norway

**Keywords:** Epigenomics, gene expression, whole-genome bisulfite sequencing, RNA sequencing, fish, hypoxia

## Abstract

Oxygen is essential for cellular metabolism, and its absence poses a severe risk to vertebrates. However, the crucian carp (*Carassius carassius*) can survive without oxygen for months due to physiological adaptations. This study investigates the role of epigenomic regulation, specifically DNA methylation, in response to anoxia and subsequent reoxygenation. Brain samples from crucian carp subjected to normoxia, anoxia, and reoxygenation (n = 4 per condition) were analyzed through transcriptome (mRNA) and whole-genome bisulfite sequencing. Results revealed a notable transcriptional response to anoxia, with 1,265 differentially expressed genes (DEGs) identified compared to normoxia. Using the machine-learning-based MethylScore, we found 14 genes with differentially methylated regions (DMRs) during anoxia. These DMR-associated genes did not meet the strict DEGs criteria (adjusted p-value < 0.05; absolute log2fold-change > 0.38), but 17 genes exhibited differences in mRNA abundance when focusing solely on DMRs. DMRs were primarily located within transcriptional start sites, and analysis of protein products revealed that genes involved in transcription regulation, protein trafficking, and signal transduction were differentially methylated and expressed under varying conditions. This study advances our understanding of epigenomic regulation in oxygen deprivation and highlights the potential in exploring alternative epigenomic mechanisms.

## Background

Oxygen is vital to cellular energy production in vertebrates and must be present continuously to support aerobic ATP production. Thus, in the animal kingdom, few vertebrates can sustain life in the absence of oxygen. In humans, oxygen deprivation plays a key role in ischemic conditions, such as stroke and heart infarctions, where the return of oxygen during reperfusion can further exacerbate tissue damage (Xie, Kittur, Li, & Hung, 2022). The crucian carp (*Carassius carassius*) is an astonishing fish that can survive without oxygen for several months at low temperatures (Holopainen, Hyvärinen, & Piironen, 1986; Vornanen, Stecyk, & Nilsson, 2009). Some of the events associated with ischemia-reperfusion, namely the removal and regain of access to oxygen, are also present in the annual fluctuations of the crucian carp life cycle. Crucian carp are endemic to Europe and Asia, inhabiting small ponds that become ice- and snow-covered during winter, which prevents photosynthesis and thereby oxygen production (Vornanen, Stecyk, & Nilsson, 2009; Holopainen, Tonn, & Paszkowski, 1997). As a result, the available oxygen decreases to a complete absence (anoxia) during winter (Holopainen, Tonn, & Paszkowski, 1997). Crucian carp sustain metabolism during winter by switching to anaerobic metabolism, relying on large liver glycogen stores, and depressing ATP consumption while increasing the glycolytic rate (Vornanen, Stecyk, & Nilsson, 2009). They reduce ATP needs by lowering central nervous system activity and movement (Johansson, Nilsson, & Törnblom, 1995; Lutz & Nilsson, 1997; Haverinen, Badr, Eskelinen, & Vornanen, 2024; Boutilier, 2001). Their skeletal muscles produce ethanol as a metabolic end product, thereby avoiding lactic acidosis when excreting the ethanol through their gills (Lefevre & Nilsson, 2024). Despite critically low oxygen levels, they preserve brain tissue with minimal damage during anoxia, though a slight increase in cell death is observed upon reoxygenation (Lefevre, et al., 2017). Overall, the physiological adaptations of crucian carp on a whole-organism level are already quite well understood (for a recent review, see (Lefevre & Nilsson, 2023)), but little is known about the molecular regulation of the responses. Since oxygen fluctuations occur yearly in crucian carp, the physiological and cellular responses must be temporary and reversible.

Epigenetics describes the stable inheritance of cellular identities through changes in gene expression that occur without altering the DNA sequence. Epigenomics refers to the DNA-associated physical and functional entities, such as histone modifications and DNA methylation, which are not necessarily inherited. These dynamic epigenomic features can rapidly change in response to environmental conditions and encompass both permanent alterations (e.g., tissue differentiation and development) and reversible changes (e.g., stress responses) triggered by various factors (Dupont, Armant, & Brenner, 2009; Holliday, 1987; Wijenayake & Storey, 2016; Struhl, 2024). The most recognized forms of epigenomic regulation include DNA methylation, histone modifications, and non-coding RNAs (Li, 2021). While histone modifications and chromatin openness may be more flexible, we focused on DNA methylation due to observed changes in turtles and fish related to hypoxia (Puvanendran, et al., 2023; Kelly, et al., 2020; Beemelmanns, et al., 2021; Ruhr, et al., 2021). DNA methylation is one of the most widespread and well-studied modifications in mammals. 5-methylcytosine (5mC) is the prevalent mark of DNA methylation in eukaryotes, while N6-methyladenine (6mA) and 4-methylcytosine (4mC) are more common in prokaryotes (Li, 2021). Unlike changes in the DNA sequence itself, which are stable, DNA methylation patterns are flexible and reversible (Kulis & Esteller, 2010). This enables organisms to respond and adapt rapidly to changing environmental cues. During the epigenomic process of DNA methylation, a methyl group is added to the fifth position of cytosine (Gong, et al., 2022). Often, 5mCs occur at cytosine-phosphate-guanine (CpG) dinucleotides, which themselves can be concentrated in large clusters called CpG islands (see Figure 1). Those are often enriched in the promoter and/or the first exon and are generally associated with transcriptional repression when methylated (Waechter & Baserga, 1982; Smith & Meissner, 2013; Saxonov, Berg, & Brutlag, 2006). Due to the impact of DNA methylation on the regulation of gene transcription, by directly influencing the accessibility of transcription factors and other regulatory proteins to the DNA, it can be viewed much as a “switch” mechanism. Altered DNA methylation patterns have been documented to affect gene expression in response to changing environmental factors in various organisms, including fish. The influence of temperature-modulated gene expression and DNA methylation was explored in the early life stages of Atlantic cod (*Gadus morhua*) by Puvanendran et al. (2023), while the regulation of hypoxia-response genes was examined during early development in Atlantic salmon (*Salmo salar*) (Puvanendran, et al., 2023; Kelly, et al., 2020). Another study by Beemelmanns et al. (2021) investigated whether temperature and hypoxia exposure affected the methylation of CpG sites and if those correlated with transcriptional changes in the liver of Atlantic salmon (Beemelmanns, et al., 2021). Finally, Ruhr et al. (2021) investigated the impact of hypoxia on the cardiovascular system of juvenile snapping turtles (*Chrysemys picta*), examining the role of epigenomic regulation via DNA methylation (Ruhr, et al., 2021).

**Figure 1:**
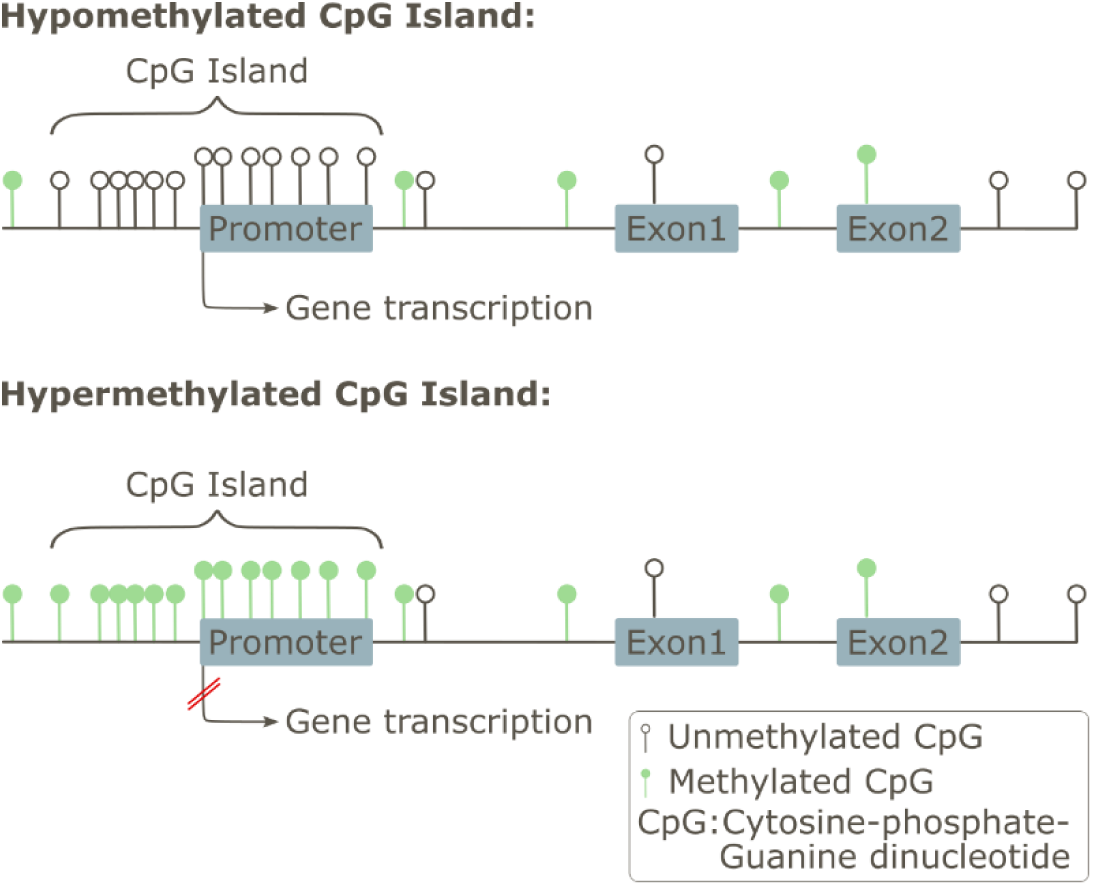
Schematic presentation of hyper- and hypo-methylation of CpG islands and the consequences for gene transcription. CpG islands are DNA regions rich in cytosine-phosphate-guanine dinucleotides located near or within gene promoter regions. When these CpG islands remain unmethylated, the associated gene is usually transcribed, whereas methylation of CpG islands typically prevents transcription.

In the present study, due to the already existing indications of a role of DNA methylation in other species, we aimed to investigate whether changes in DNA methylation patterns occur in response to anoxia and reoxygenation in the crucian carp and to what extent these changes correlate with alterations in gene expression. The remarkable ability of crucian carp to undergo such drastic changes in their physiology to survive anoxia offered a unique opportunity to explore the epigenomic mechanisms involved in the physiological and cellular response to oxygen deprivation. The brain was chosen because its high energy demand and thus sensitivity to oxygen deprivation means that the responses to anoxia may be particularly pronounced in this tissue. We utilized next-generation whole-genome bisulfite sequencing (WGBS) in combination with mRNA sequencing to investigate altered gene-environment interactions, providing comprehensive, high-resolution, and quantitative methylation data. By correlating DMR-associated genes and DEGs during the transition from normoxia to anoxia and back to normoxia through reoxygenation, we aimed to identify key genes involved in the response to anoxia. This study advances our understanding of the physiological responses of the crucian carp brain under oxygen stress conditions and adds additional information to the functional annotation of the crucian carp genome.

## Material and Methods

### Crucian Carp Husbandry and Exposure Experiment

Crucian carp involved in this study (total N=12, n=4 per condition, both sexes, mean mass = 46.26 ± 1.80 *g*) were obtained from the Tjernsrudtjernet pond (Oslo, Norway, N59.92151, E10.60973) in September 2020. The fish were collected during autumn to ensure their readiness for cold winter conditions when they usually experience anoxia in nature (Vornanen, Stecyk, & Nilsson, 2009). The animals were kept in a 500-liter holding tank in the aquarium facilities at the Department of Biosciences, University of Oslo. The fish were subjected to a 12 h darkness / 12 h daylight cycle and supplied with aerated and dechlorinated Oslo tab water at 9 – 10°C. The animals were fed daily with commercial carp food for approximately four months and maintained under stable conditions to acclimate them to the facility and assess their health and feeding behavior. The water quality was checked once a week with Tetra test kits for pH, ammonia/ammonium (NH3/NH4^+^), nitrite (NO2^-^), and nitrate (NO3^-^). Removing the animals from their natural habitat and acclimatizing them to the animal facility and the subsequent experiment were approved by the Norwegian Food Safety Authority (FOTS ID 16063). To mimic natural conditions, the fish were not fed during exposure and the 24-hour acclimatization. Anoxic fish, with reduced physical activity, are unlikely to eat, and feeding them could introduce uncontrolled variables and degrade water quality (Holopainen, Hyvärinen, & Piironen, 1986; Nilsson, 1990). Three identical 25-liter cylindrical dark buckets, each featuring a 5 x 10 cm clear plastic window covered with plastic sheets for daily health checks, were equipped with aerated 7°C water in a flow-through system. One bucket, continuously supplied with aerated water, served as the normoxic control, while the others were completely (anoxia) or temporarily (reoxygenation) deprived of the aerated water supply. Thirty fish were placed in each bucket, with four from each condition sampled for this study, while the remaining fish were used for other purposes within the research group. There was no mortality. An equal number of fish in each tank ensured that the environmental conditions were the same in all buckets except for the oxygen supply. Oxygen saturation in each bucket and the ambient water temperature were monitored daily with a FireSting-O2 (4 channels, PyroScience GmbH). During exposure, 90% air saturation was maintained by bubbling air in the normoxic bucket. Air bubbling was replaced with nitrogen gas in the anoxic and reoxygenation buckets during the anoxic period. Both buckets maintained their oxygen levels below the detection limit of the oxygen meter and hence were considered anoxic. After the anoxic period, reoxygenation of the water was restored by discontinuing the nitrogen gas supply and adding air bubbling in the reoxygenation bucket. The normoxic (n=4) and anoxic (n=4) groups were sampled on the seventh day, resulting in a five-day-long exposure to anoxia. The remaining reoxygenation (n=4) group received 24 hours of reoxygenation and was sampled the day after. For the omics analysis, we utilized four biological replicates from each condition bucket, successfully including both male and female fish in one condition. However, due to the difficulty of determining sex before sampling, we could not guarantee equal representation of both sexes in each condition.

### Tissue Sampling and Crushing

During sampling, the fish were netted one at a time and immediately euthanized with a sharp blow to the head, followed by cervical transection behind the head, adhering to the only approved method for euthanizing fish in Norway. The brain tissue was extracted within a minute. An anteriorly oriented cut from the incision towards the eyes opened the cranium, severing the olfactory tract, optic nerve, and spinal cord, allowing the extraction of the entire brain tissue in one piece. The brain tissue was immediately frozen in liquid nitrogen and stored at -80°C. Each brain (mean mass = 40.45 ± 12.00 *mg*) was crushed with the BioPulverizer (BIOSPECPRODUCTS) to ensure uniform distribution of all brain components in the sample, and the tissue powder was split into two aliquots. Subsequently, DNA and RNA were extracted as described below (see section *DNA Isolation, Library Preparation, and Sequencing* and *RNA Isolation, Library Preparation, and Sequencing*). Given the challenge of consistently extracting good qualitative DNA and RNA using a single method (utilizing the AllPrep DNA/RNA Mini Kit (from QIAGEN), we adopted a dual-method approach to obtain high-quality extracts for both DNA and RNA.

### DNA Isolation, Library Preparation, and Sequencing

DNA extraction from crushed brain tissue was accomplished using the AllPrep DNA/RNA Mini Kit (QIAGEN). Tissue powder was combined with 3 mm Tungsten Carbide Beads, submerged in lysis buffer, and then homogenized using the Tissue Lyser II (QIAGEN). The protocol provided by the manufacturer was followed without modifications. The quality of DNA was evaluated using the NanoDrop 2000 Spectrophotometer (ThermoFisher Scientific). Analysis of the DNA samples revealed an average yield of 538.54 ± 238.541 ng µl^-1^ and a A260 / 280 value of 1.85 ± 0.01 and a A260 / 230 value of 2.10 ± 0.42, indicating high purity as described by (Lucena-Aguilar, et al., 2016). The samples were kept at -20°C until the library preparation.

The Pico Methyl-Seq^TM^ Library Prep Kit (Zymo Research) was used for genome-wide DNA methylation profiling. This technique involves library preparation, amplification, and quality control (Gong, et al., 2022). Starting the protocol with sample concentrations of 2.5 ng µl^-1^, the DNA conversion by bisulfite treatment was followed by adapter attachment to the fragment extremities. During the bisulfite transformation, unmethylated cysteines are converted to uracil, while methylated cytosine residues remain unchanged (Gong, et al., 2022). Modifications were made to the amplification stage of the protocol as the Zymo-Seq UDI Primer Set (Zymo Research) was used instead of the Index Primer provided by the kit. The solution quantities were adjusted to match the modified indexes, and six amplification cycles were performed. The DNA fragments were purified according to the DNA Clean & Concentrator^TM^ MagBead Kit manual. A left-sided selection for fragments >100 bp was performed with a MagBeads ratio of 1.24. Afterward, the concentration of the product was determined with the Qubit 3 Fluorometer (Invitrogen, ThermoFisher Scientific), and the quality of the library was assessed with the Bioanalyzer High Sensitivity DNA Analysis Chips (Agilent). The converted DNA was sequenced at the Norwegian Sequencing Center (one NovaSeq 6000 SP flowcell and two quarters of a NovaSeq 6000 S4 flowcell). Our study included three conditions with four replicates per group (Ntotal=12) and a per-sample coverage of around 22 times, exceeding the required range of 5 – 15 times, as advised by (Ziller, Hansen, Meissner, & Aryee, 2014).

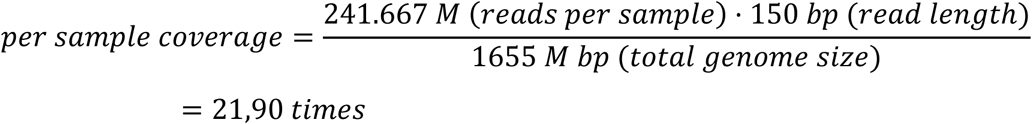

### RNA Isolation, Library Preparation, and Sequencing

RNA extraction from crushed brain tissue was conducted using the TRIzol Reagent (ThermoFisher Scientific). Upon adding Trizol and Tungsten Carbide Beads (3 mm) to the tissue powder, the samples were homogenized for one minute at a frequency of 30 Hz in the Tissue Lyser II (QIAGEN). Subsequently, the aqueous phase was separated, and 10 µl of RNAse-free glycogen was supplemented to enhance yield. Following precipitation and washing, RNA solubilization was carried out in 30 – 40 µl of RNAse-free water. The quality of extracted RNA was assessed using the NanoDrop 2000 Spectrophotometer (ThermoFisher Scientific). The RNA samples demonstrated an average yield of 751.29 ± 350.63 ng µl^-1^ and a A260 / 280 ratio of 1.88 ± 0.08 and a A260 / 230 value of 1.33 ± 0.28, indicating good purity. RIN numbers, not descending below 8.6 and exhibiting an average value of 9.15 ± 0.32, confirmed high RNA integrity. Subsequently, the Norwegian Sequencing Center performed library preparation using the TruSeq® Stranded mRNA Library Prep kit (Illumina) and sequencing on a half Illumina NovaSeq SP, yielding an average of 16.7 M reads per sample.

### Bioinformatical Analysis

Unless otherwise stated, computations were done on the high-performance computing cluster “Saga” provided by Sigma2 - the National Infrastructure for High-Performance Computing and Data Storage in Norway. DNA and RNA sequencing datasets were initially assessed for quality using FASTQC (v0.11.9-Java-11). Following this, adapter clipping and quality trimming were performed using TrimGalore (v0.6.10-GCCcore-11.2.0). For DNA trimming, we applied the settings “--paired --quality 20 --length 20 --clip_R1 10 --clip_R2 10 --three_prime_clip_R1 10 --three_prime_clip_R2 10”. The settings “--paired --quality 20” were used for RNA trimming. Trimmed reads were subsequently mapped to the crucian carp genome using BWA-METH (v0.2.7; default settings) for DNA sequences and HISAT2 (v2.2.1- gompi-2022a) with the settings “--reorder --summary-file” for RNA sequences. The genome assembly and annotation were sourced from Valencia-Pesqueira et al. (Valencia-Pesqueira L. M., Hoff, Tørresen, Jentoft, & Lefevre, 2025). Note that the annotation was only used to map RNA reads. Mapping results were converted to BAM files using SAMtools (v1.17-GCC-12.2.0). The featureCounts module of Subread (v2.0.3-GCC-11.2.0) was used with the settings “-T 4 - O -C -p -s 2 -t exon -g gene_id” to generate exon counts summarized at the gene level from the RNA sequencing data, allowing multi-overlapping reads, but discarding chimeric alignments. FeatureCounts excluded multimapped reads from its analysis, ensuring that only uniquely mapped read counts were considered (for raw output from featureCounts, refer to Supplementary Material 01). MethylScore (v0.2; https://github.com/Computomics/MethylScore) was employed to identify DMRs with the following parameters: DMR_MIN_C=5 (minimum five cytosines in each DMR), DMR_MIN_COV=3X (minimum three times coverage in each cytosine), MR_FREQ_CHANGE=20 (at least 20% of samples showing a change in a methylated region - frequency to be tested as a candidate DMR), CLUSTER_MIN_METH_DIFF=20 (which sets a 20% cutoff for methylation difference between clusters in CG, CHG, and CHH contexts) (Hüther, et al., 2022). All other parameters used the default settings. We selected MethylScore for its ability to handle genome duplication, as it was initially developed for complex plant genomes, such as *Arabidopsis thaliana*, making it suitable for the duplicated genome of crucian carp (Valencia-Pesqueira L. M., Hoff, Tørresen, Jentoft, & Lefevre, 2025). For the DMR identification with MethylScore, we performed comparisons across conditions: N to A, N to R, and A to R. The following analyses were done in Python. The initial clustering output was filtered to retain only clusters consistent with a single experimental condition. A cluster was kept only if it contained samples exclusively from one condition (e.g., solely normoxic samples). To ensure robust coverage, we filtered the DMRs to include only those represented by at least six biological samples, meaning a minimum of three biological samples from each of the two compared conditions or four biological samples from one condition and two biological samples from the other tested condition. This criterion ensured that no single sample disproportionately influenced the identification of the DMR.

Since the used annotation lacked information on promoter regions and untranslated regions (UTRs) are included for most genes but not all, we extended our analysis by adding five percent of the gene length upstream of the first exon (in example below: the RBS region) and downstream of the last exon (in example below: the termination region) of the coding domain sequence, which we now refer to as the "gene borders" (see Figure 2). Gene models (i.e. the annotation.gtf file), including a five percent buffer for UTRs and promoter regions, were used for gene-to-DMR mapping.

**Figure 2:**
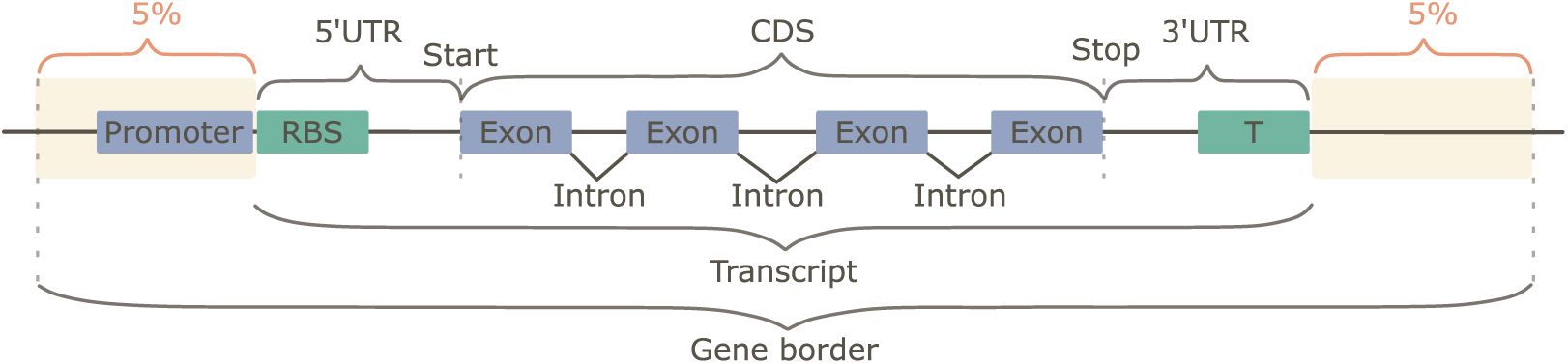
Schematic representation of the "gene border" which includes a promoter that initiates transcription. The transcript entails the ribosome binding site (RBS) in the 5’UTR region, the translation start site (Start), the coding domain sequence (CDS), the translation stop site (Stop), and the 3’ UTR region ending with the terminator (T). While the existing gene annotation accounts for most UTR regions, a 5% margin (orange box) was added to address omissions and promoter regions. This 5% margin will typically exclude UTRs when they are already annotated. However, for some genes lacking UTR annotations, the 5% margin will encompass these regions. The utilized annotation includes 45,667 genes, of which 44,666 are annotated with 5’ UTRs and 45,265 with 3’ UTR regions. Collectively, this is referred to as the gene border.

For the examination of the distribution of DMRs within the gene borders, we categorized them into six regions: ‘Before Start’ (upstream of transcription start), ‘Overlapping Start’ (spanning from the untranscribed region into the transcribed region, overlapping the transcriptional start), ‘Inside Gene’ (entirely within the transcribed region of a gene), ‘Overlapping Stop’ (spanning from the transcribed region into the downstream untranscribed region, overlapping the transcriptional stop), ‘After Stop’ (downstream of transcriptional stop including the untranscribed region), and ‘Overlapping Both’ (overlapping transcriptional start and stop). The methylation status, defined as hypermethylated or hypomethylated, was determined by comparing average methylation between conditions. The average methylation rate for each cluster, as reported by MethylScore, was further averaged across all clusters within the same biological condition (e.g. normoxia, anoxia). These condition-specific averages were then compared to determine hypermethylation (e.g. normoxia < anoxia) or hypomethylation (e.g. normoxia > anoxia) relative to normoxia.

### Statistics

Differentially expressed genes were detected using the DESeq2 (v1.40.2) package in R. DEGs were identified in Python after filtering out rows with fewer than ten gene counts, applying an absolute log2(fold-change) threshold of > 0.38 (corresponding to an increase or decrease of 30%, i.e. fold-change larger than 1.3 or smaller than 0.77), and selecting only those with an adjusted p-value of <0.05 for further analysis. Gene Ontology (GO) enrichment analysis was conducted using the “goseq” package (v1.52.0) in R, and redundant GO terms were removed using Revigo (http://revigo.irb.hr/) (Supek, Bošnjak, Škunca, & Šmuc, 2011). GO terms were extracted for genes based on corresponding UniProtKB/Swiss-Prot proteins. Of 45,667 genes, 41,373 had a Swiss-Prot protein match and associated GO term(s) (Valencia-Pesqueira L. M., Hoff, Tørresen, Jentoft, & Lefevre, 2025). P-values adjusted for multiple testing were obtained through a Bonferroni correction of the p-values computed by goseq. The negative log-transformed adjusted p-values were estimated to facilitate the identification of statistically significant enrichments. The expected number of DEGs in each GO category was calculated as the number of DEGs one would expect to find in that GO category, based on the proportional representation of genes in that category in the complete gene set. Fold enrichment was calculated by dividing the observed number of DEGs within each GO category by the expected number of DEGs in that category, thereby estimating the increased prevalence of DEGs relative to the background gene set.

Differences in mRNA abundance of DMR genes were assessed in Python by first normalizing the data using the trimmed mean of M-values (TMM) method (Robinson & Oshlack, 2010). When the standard deviation was within 0.5 of the mean standard deviation, a one-way ANOVA test was used to test for significance. Otherwise, the Alexander-Govern test was applied. A Tukey post-hoc test was conducted in both cases to identify significant differences in pairwise comparisons. For heatmap visualization of genes with significant differences in methylation pattern (DMR genes), we filtered the expression data (mRNA abundance) to include a minimum of 50 counts across all samples and normalized the data using z-score scaling. All plotting was done in Python.

## Results

We investigated changes in DNA methylation and gene expression under normoxia (N), anoxia (A), and reoxygenation (R) conditions in crucian carp brain tissue using WGBS and mRNA sequencing. The WGBS data had an average mapping efficiency of 95% uniquely aligned reads with BWA-METH, while the RNA data achieved 96% uniquely aligned reads with HISAT2. The principal component analysis (PCA) of the mRNA sequencing data confirmed that the 12 biological replicates clustered according to the three experimental conditions. Furthermore, the inclusion of the single normoxic female within an otherwise tightly clustered, normoxic group of males suggests an absence of sex-specific differences (Figure 3A). Notably, the samples in the anoxia and reoxygenation conditions were more dispersed than the tight clustering observed in the normoxia condition, indicating greater variability in gene expression with anoxia treatment than between males and females. We investigated differential gene expression across the three comparisons [normoxia to anoxia (N to A), normoxia to reoxygenation (N to R), and anoxia to reoxygenation (A to R)] and identified hundreds of DEGs (Figure 3B, C; for raw output from the DESeq2 analysis, refer to Supplementary Material 02 - 04). In the A to R comparison, we found the fewest DEGs, 145 downregulated and 317 upregulated. The N to A comparison showed the highest number of genes with altered mRNA abundance, with 720 upregulated and 545 downregulated genes. The N to R comparison identified 394 upregulated and 186 downregulated genes. Next, we examined whether genes overlapped across comparisons and grouped DEGs accordingly. Of the 1265 DEGs in the N to A comparison, 306 were still differentially expressed in reoxygenation. Meanwhile, 189 genes differentially expressed in anoxia had changed significantly from anoxia to reoxygenation but were no longer differentially expressed compared to normoxia. There were 252 genes differentially expressed in reoxygenation, of which 183 were specific to reoxygenation, and 69 were different also in anoxia (Figure 3B). Out of the 45,667 annotated genes (Figure 3D), 40,259 had a minimum coverage of ten reads and were thus included in the differential expression analysis.

**Figure 3:**
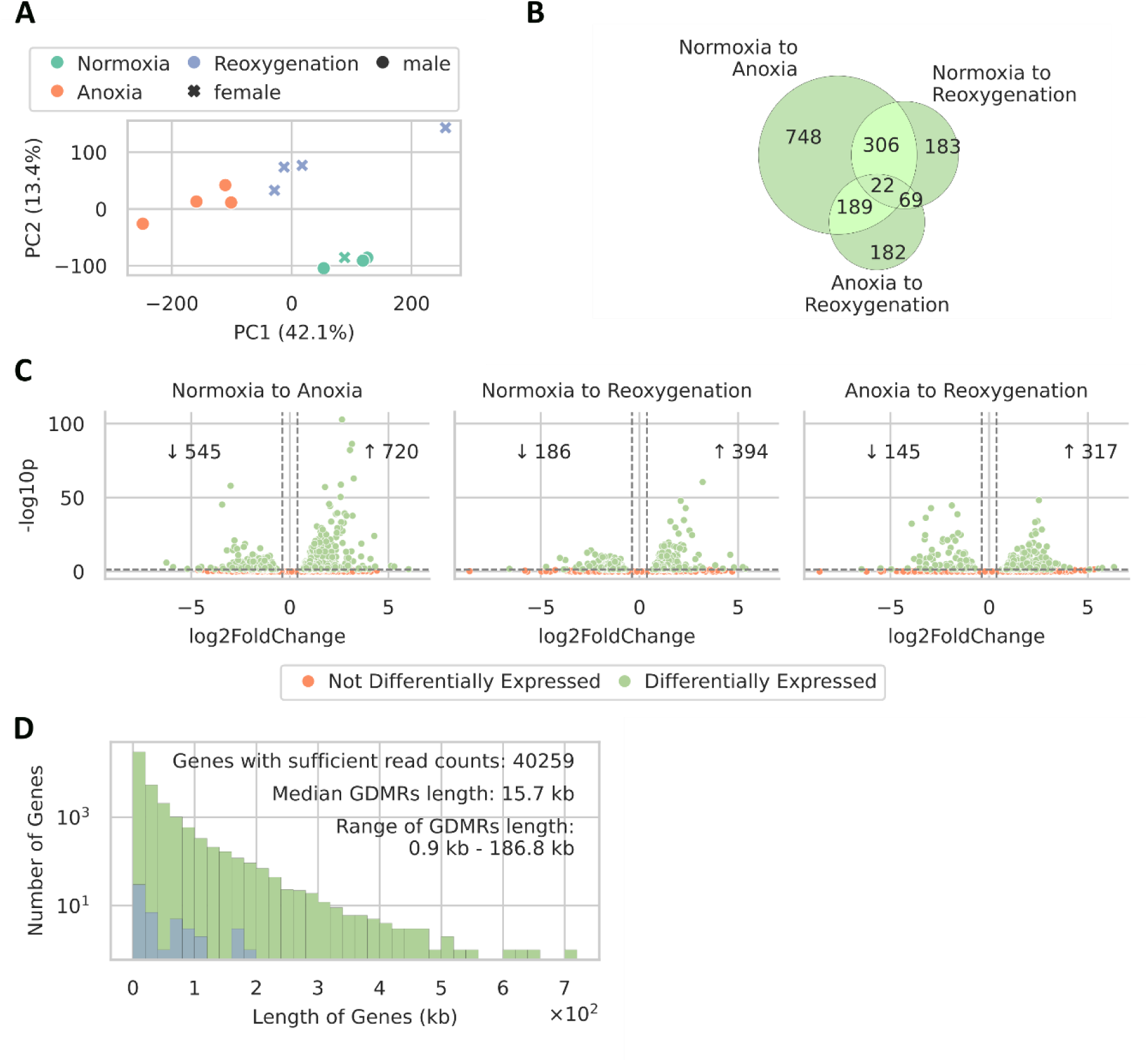
Transcriptome of the different conditions and comparisons. A: PCA analysis of sample clustering in the mRNA sequencing data in the four biological replicates for the three conditions. B: Venn diagram of common differentially expressed genes between the comparisons. C: Significantly differentially expressed genes (DEGs). The plot highlights DEGs between the treatment conditions, using p-value < 0.05 and absolute log2 fold change ≥ 0.38. Insets show the number of genes with increased or decreased expression in each comparison. Upregulation refers to higher abundance in the second condition compared to the first. D: Length distribution of annotated protein-coding genes (green) and DMR genes (GDMRs; blue) in the genome. Lengths are in kilobases (kb). For the raw output from the DESeq2 analysis, please see Supplementary Material 02 – 04.

We further conducted a GO enrichment analysis to elucidate putative biological functions and implications of DEGs, regardless of the direction of change in abundance (Figure 4; for raw GO enrichment analysis output, refer to Supplementary Material 05 - 07). Given our interest in the transition to and endurance of anoxia, we also performed a GO enrichment analysis specifically for the N to A comparison, categorizing up- and down-regulated genes in the same row format (Figure 5). Whether we analyzed upregulated or downregulated genes in the N to A comparison, or examined the N to R and A to R comparisons, the identified enriched GO terms remained specific and involved only a few genes (Figure 4 and Figure 5). However, an exception was found in the row corresponding to the cellular component (CC) terms in the N to A comparison, where the term "nucleus" was associated with either 349 or 232 genes. Given the general nature of the "nucleus" term, it is unsurprising that it encompasses many genes. Only the most significant terms (adjusted p-value < 0.05) are displayed, with the highest significance in the A to R comparison.

**Figure 4:**
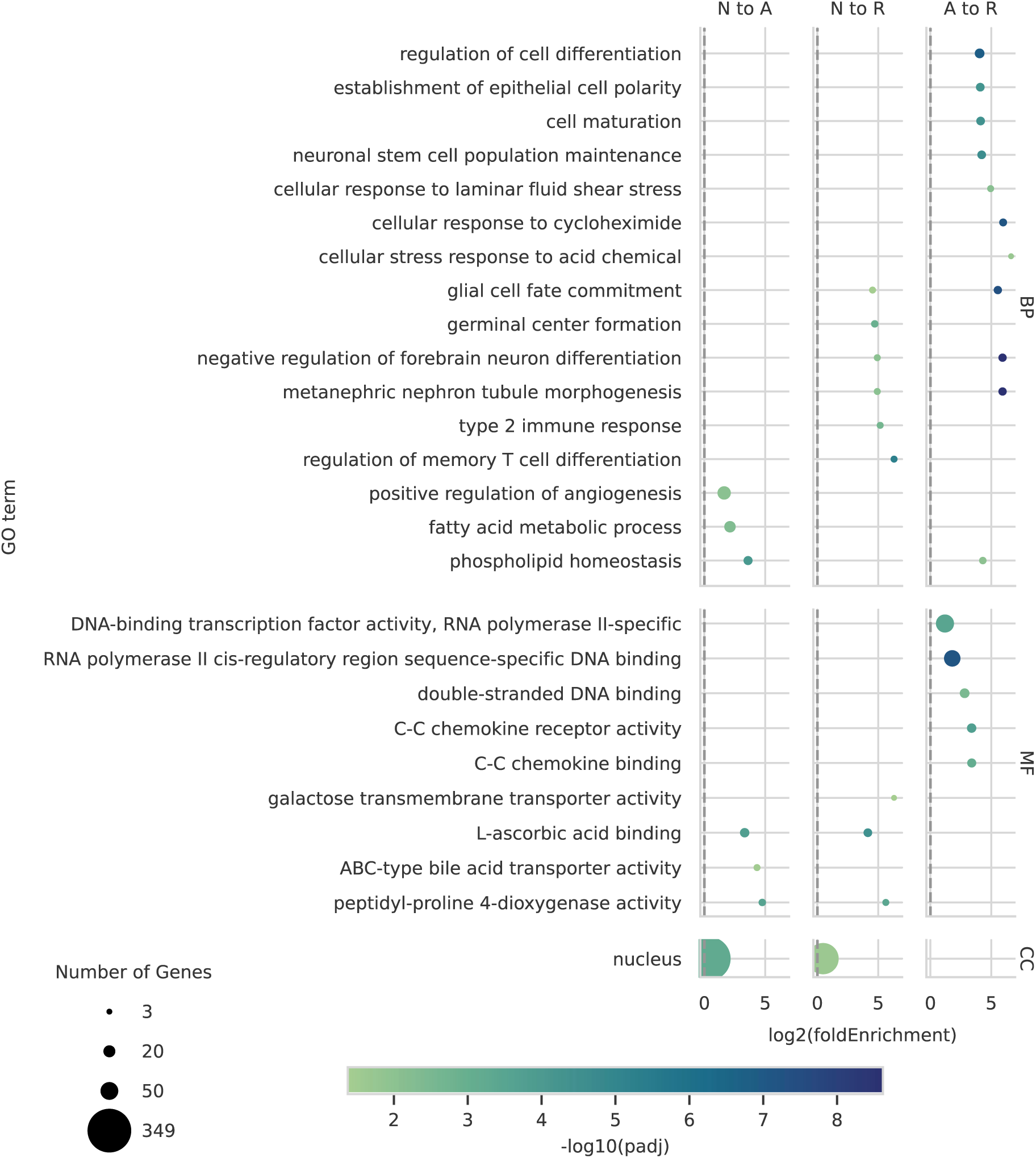
Gene ontology enrichment analysis of the three comparisons (N to A, N to R, A to R). The plot is structured into three columns for each comparison and separate rows for gene ontology terms in biological processes (BP), molecular functions (MF), and cellular components (CC). Log2(fold enrichment) indicates how many more times a given term appears amongst the DEGs than what is expected based on the frequency in the entire set of genes. For the raw GO enrichment analysis output, please see Supplementary Material 05 – 07.

**Figure 5:**
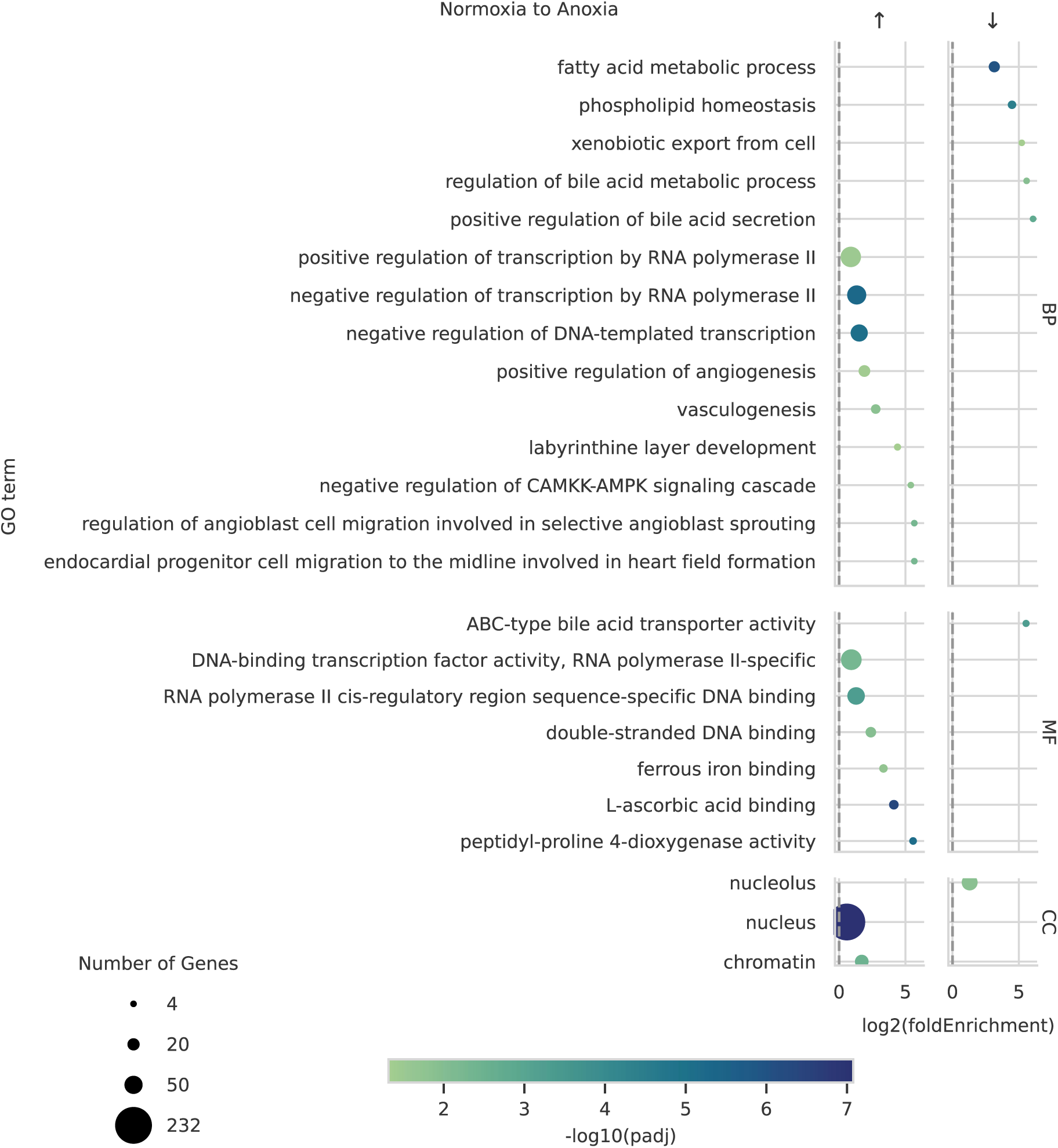
Gene ontology enrichment analysis comparing normoxia to anoxia. shows upregulated terms on the left panel and downregulated terms on the right. The plot is structured into separate rows for gene ontology terms in biological processes (BP), molecular functions (MF), and cellular components (CC). Log2(fold enrichment) indicates how many more times a given term appears amongst the DEGs than what is expected based on the frequency in the entire set of genes. For the raw GO enrichment analysis output, please see Supplementary Material 07.

From the DNA sequencing data, we identified a total of 53 genes with DMRs located within their gene borders across the three comparisons (Figure 6A; for the raw MethylScore DMR identification output, see Supplementary Materials 08-10). For N to A, we identified 14 genes with DMRs within the gene borders, 28 genes for N to R, and 15 genes for A to R. The median DMR gene length was 15.7 kb, ranging from 0.9 to 186.6 kb. The length distributions of mRNA transcripts and DMR genes were comparable (Figure 3B), indicating that detection of DMRs was not biased towards long genes. Furthermore, we investigated the overlap of DMR-regulated genes across different comparisons (Figure 6B). Two genes were shared between the N to A and N to R comparisons, indicating that these two genes were DMR in anoxia as well as reoxygenation but not different from A to R. Another two were shared between the N to A and A to R comparisons, indicating that the genes were DMR in anoxia, and had changed pattern in reoxygenation relative to anoxia but no longer different from normoxia (Figure 6B). Next, we examined the distribution of DMRs within the gene borders (Figure 6C, F). In all three comparisons, most DMRs were inside the gene, i.e. in the transcribed region. There were a few occurrences of DMRs before the transcriptional start site, but even more were found overlapping the transcriptional start site, with the majority of DMRs located within the gene itself (transcribed region). Minor occurrences of DMRs were observed overlapping the transcriptional stop (one ‘Overlapping Stop’ in N to R and A to R) and after the transcriptional stop (two ‘After Stop’ in each comparison). No DMRs were found to overlap both the start and stop regions. Furthermore, we investigated whether genes contained one or multiple DMRs (Figure 6D). In the N to A comparison, 13.3 % of genes featured two DMRs, whereas in all other comparisons, genes consistently included only a single DMR. We utilized a staircase plot to examine the length and coverage of DMRs within genes (Figure 6E). For the N to A comparison, DMRs showed minimal overlap and were generally short, occurring evenly distributed before the transcriptional start, within the gene, or after the stop. In the N to R comparison, many DMRs cluster within the first 25% of gene length with a significant overlap of up to six DMRs, followed by a long stretch without DMRs and some clustering at the transcriptional end. In contrast, the A to R comparison featured much longer DMRs, with one notable DMR spanning from approximately 50% to 90% of the gene’s length. Although DMRs overlapped less in this comparison, they were much longer. There was no direct overlap when comparing the list of DMR genes to those classified as DEG (Figure 6G). In other words, the genes associated with DMRs were not detected to be differentially expressed with the statistical method, correction for multiple comparisons, and cut-offs applied. To gain a deeper understanding of DNA methylation dynamics, we examined the methylation status for each gene associated with DMRs within each comparison to identify regions that gain or lose DNA methylation (Figure 6H). Seven of the 14 identified DMR genes in the N to A comparison exhibited hypermethylation, while the remaining seven showed hypomethylation. In contrast, the other two comparisons showed more genes with hypomethylation, suggesting that the transition to anoxia involves a balance of gene activation and inactivation, while the transition from anoxia and back to normoxia is predominantly characterized by gene activation.

**Figure 6:**
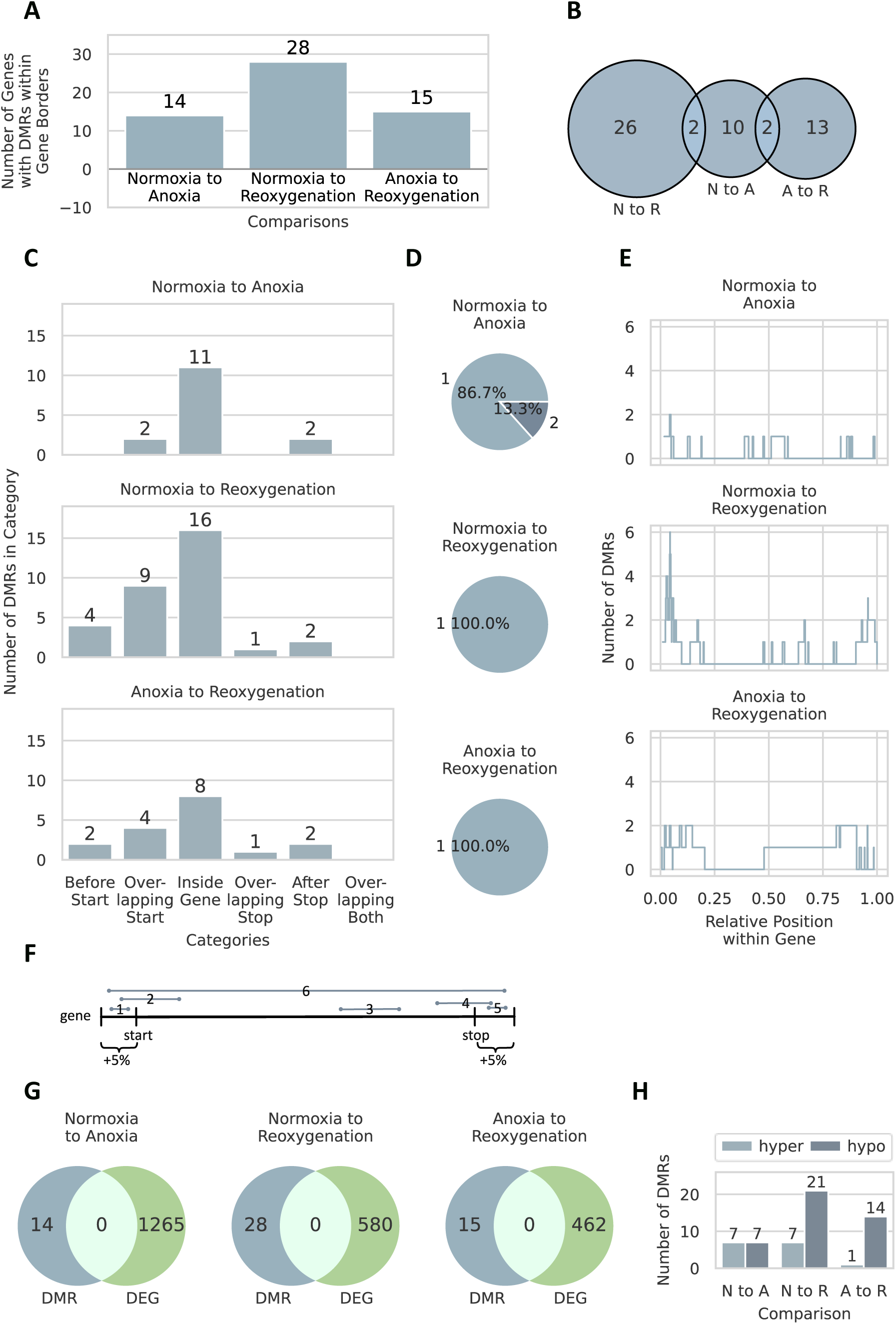
Differentially methylated regions (DMRs). A: Identified DMRs within gene borders in each comparison. B: Overlapping DMR genes between the comparisons. C: DMR locations within gene borders (for more information regarding the categories, see methods sections; for a schematic overview of the categories, see F). D: Number of DMRs per gene. E: Staircase plot of DMR coverage. This plot begins at zero on the y-axis, stepping up by one unit with the start of each DMR and descending by one unit when a DMR ends. Overlapping DMRs result in additional upward steps, with the y-axis indicating the number of overlaps and the x-axis representing the genomic length covered. F: Schematic illustration of DMR location categories (1: before start, 2: overlapping start, 3: inside gene, 4: overlapping stop, 5: after stop, 6: overlapping both; for more information, see methods section). G: Correlation of DMRs with DEGs. H: Methylation changes between comparisons. For the raw MethylScore DMR identification output, please see Supplementary Material 08 – 10.

Although no direct overlap was found between DMR genes and DEGs, we examined the mRNA abundance levels of DMR genes with sufficient counts, because the DEG analysis could have excluded genes with smaller or more variable abundance patterns. The heatmap illustrates the expression changes across different conditions and comparisons (Figure 7). Of the 53 genes with DMRs located within the gene borders, 50 had sufficient counts (the sum of the expression levels across all samples for each DMR gene ≥ 50), and 17 were significantly differentially expressed in the one-sided ANOVA or Alexander-Govern test. Normoxia samples clustered together, while anoxia and reoxygenation samples showed partial clustering, mirroring the PCA analysis of the transcriptome. The methylation status (i.e. overall hyper- or hypomethylated in one condition compared to another) of the 17 DMR genes with altered expression revealed a consistent pattern for four genes possessing DMRs adjacent to the transcriptional start site (Figure 8). In these cases, increased methylation correlated with reduced mRNA abundance, and decreased methylation coincided with increased mRNA abundance. The gene putatively coding for SPAT2 (spermatogenesis-associated protein 2, ccar_ua02-g2350) exhibited DMRs across the comparisons N to A and A to R (Figure 8A) with low methylation and upregulated expression in normoxia compared to increased methylation and downregulated expression in anoxia. Notably, SPAT2 is particularly interesting as it has DMRs in two comparisons, covering all conditions. For the gene putatively coding for CLMN (calmin), there was a DMR only from N to R, but a consistent pattern of decreasing methylation and increasing mRNA abundance was observed over exposure time (Figure 8B). The gene putatively coding for ERF (ETS domain-containing transcription factor) had a DMR from N to A. It showed approximately 10% methylation in normoxia, increased to 50% in anoxia, with decreased mRNA abundance; however, methylation levels were maintained in reoxygenation, despite increased mRNA levels (Figure 8C). SRS1B (serine/arginine-rich splicing factor 1B) showed decreasing methylation from normoxia to reoxygenation, with increased mRNA in anoxia compared to normoxia. Despite decreased methylation, mRNA levels were reduced during reoxygenation (Figure 8D). The detailed expression patterns for each of the 17 significant genes are provided in Figure 9 (listing of identified protein products see Supplementary Materials 12). Most of the 17 genes exhibited decreased mRNA abundance under anoxic conditions, except for CLMN and SRS1B (Figure 8). Although some genes returned to normoxic levels (Figure 9: B, D, E, G, H, J, K, L, M), not all fully recovered (Figure 9: A, C, F, I).

**Figure 7:**
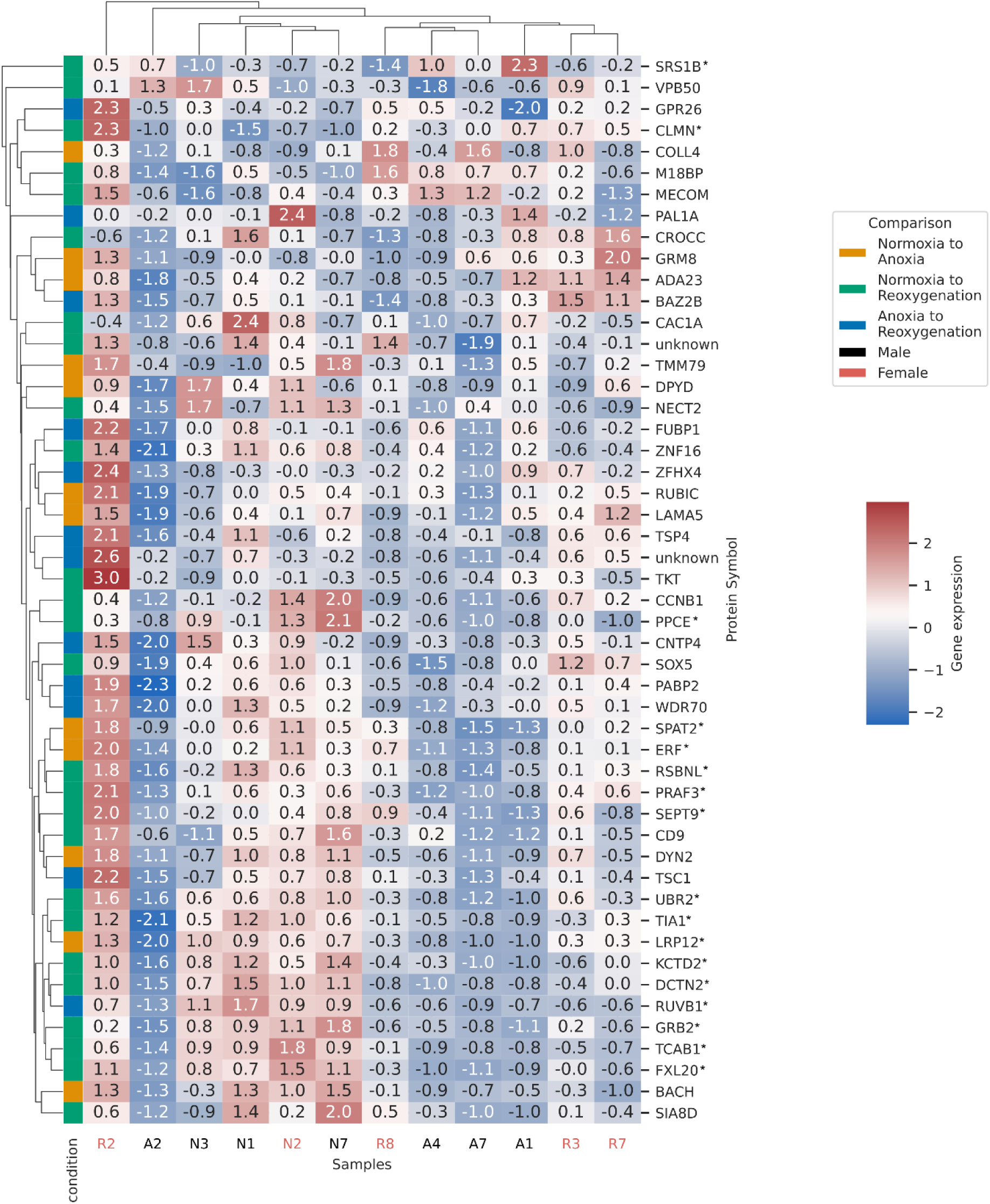
Heatmap of mRNA expression data for all DMR-related genes with sufficient coverage for all comparisons. The condition color code indicates in which treatment comparison a gene was determined to be differentially methylated. Data are scaled using the z-scale method. * indicates significant expression changes. For TMM-normalized counts, see Supplementary Material 11.

**Figure 8:**
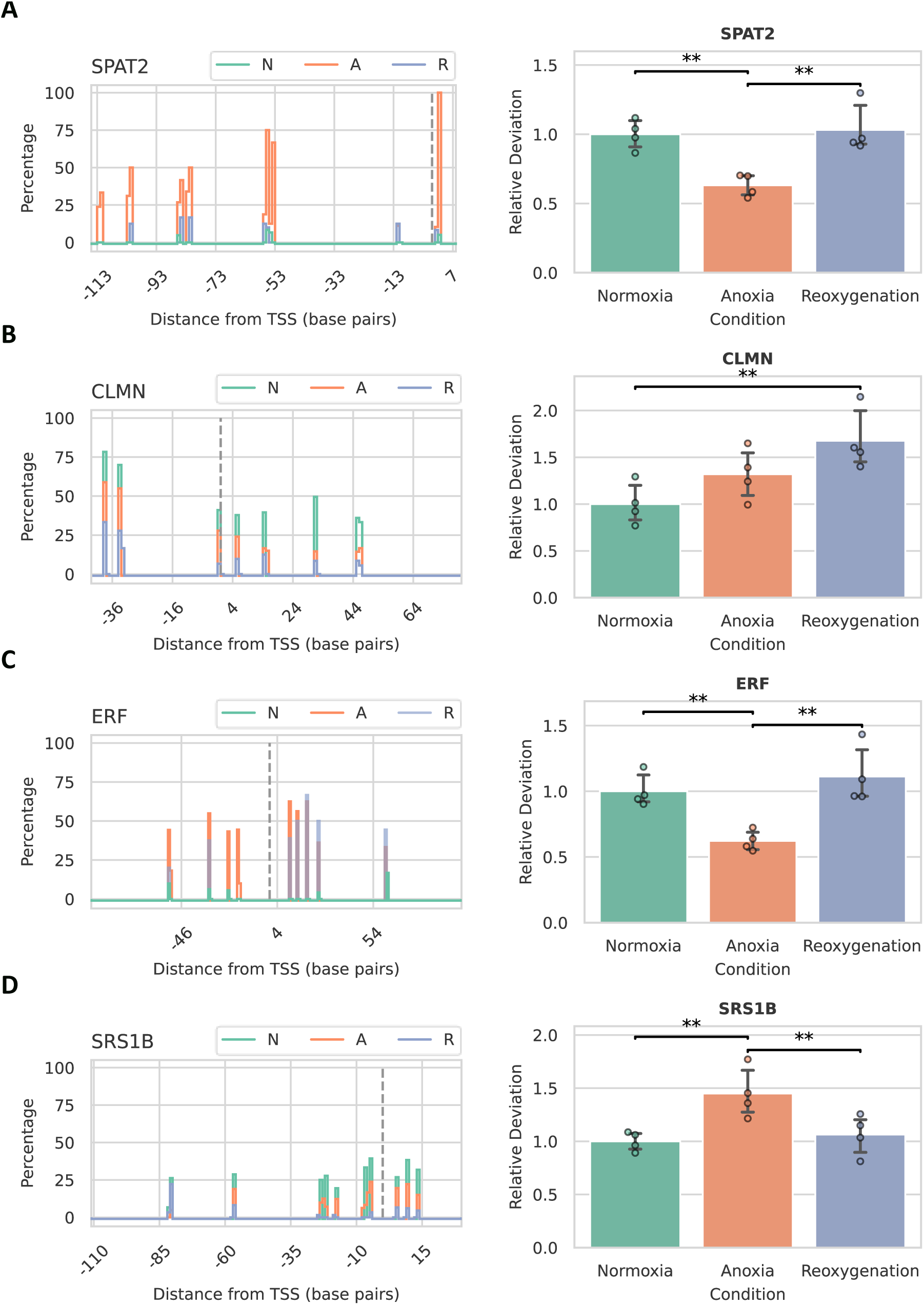
Methylation status changes of the DMR on A: SPAT2, B: CLMN, C: ERF, and D: SRS1B across three conditions: normoxia (N), anoxia (A), and reoxygenation (R). Left: Stair plot displaying average methylation sites within a DMR. Each line represents the average of all biological samples in a condition, including both forward and reverse strands. The y-axis baseline is -1, ascending by the average methylation percentage at the start of a site and descending when it ends. The y-axis indicates methylation percentage, and the x-axis represents distance from the transcription start site (TSS) (in bp). The grey dotted line represents the TSS. B: Normalized DMR gene expression data. Significance level **: 2 σ, ***: 3 σ. For TMM-normalized counts, see Supplementary Material 11.

**Figure 9:**
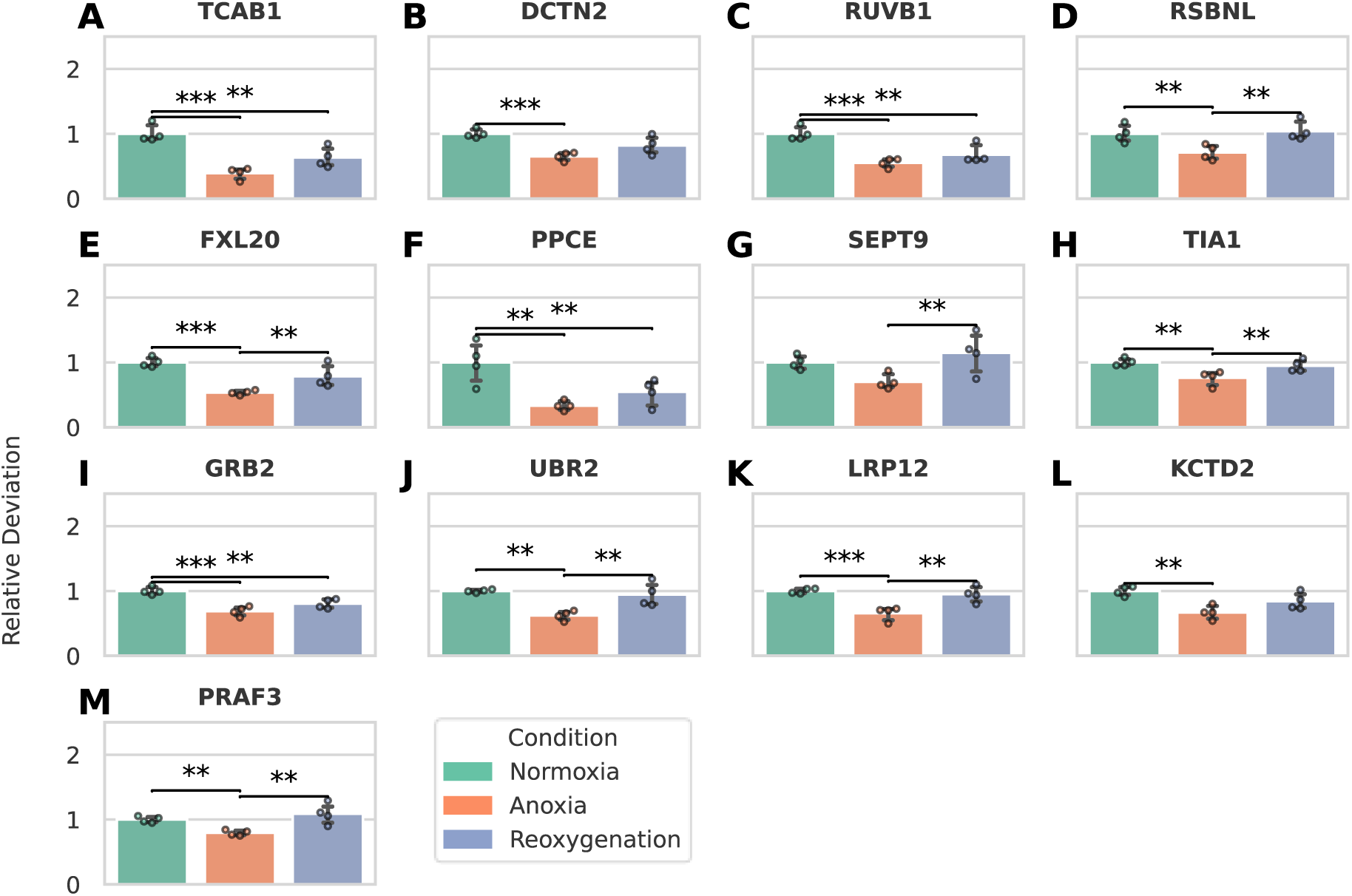
Normalized DMR gene expression data highlighting 13 of the 17 genes with altered expression (the remaining four are detailed in Figure 8) from the total DMR gene set presented in Figure 7. Significance level **: 2 σ, ***: 3 σ. For TMM-normalized counts, see Supplementary Material 11.

## Discussion

In this study, we aimed to investigate whether changes in DNA methylation patterns occur in response to anoxia and reoxygenation in the brain of crucian carp, and to what extent these changes correlate with alterations in gene expression. We found that specific genes are indeed differentially methylated in response to anoxia and reoxygenation; however, the number of differentially transcribed genes vastly exceeds the number of DMR-associated genes, indicating that other mechanisms of regulation are also at play. This study enhances our understanding of epigenomic regulation by demonstrating that differential methylation does not necessarily correlate broadly with changes in gene expression. When the abundance of DMR-associated genes was investigated more closely, it was revealed that roughly half of the genes did in fact exhibit changes in abundance, but that patterns were complex, i.e. it was not as simple as hypomethylation always correlating with increased mRNA abundance and *vice versa*. These findings emphasize that other epigenomic modifications may be involved, and we should consider additional factors such as gene context, chromatin state, and the influence of non-coding regulatory elements in a context-dependent manner.

DEG analysis of 40,259 genes meeting read count requirements highlighted key transcriptomic responses to different oxygenation states, consistent with unpublished findings from our group (Lefevre & Nilsson, 2024). The N to A comparison exhibited most changes in mRNA abundance (720 upregulated, 545 downregulated), while the A to R transition had the fewest (317 upregulated, 145 downregulated). Our findings are consistent with other studies on temperature and hypoxia stress, which also report a similar number of upregulated and downregulated genes (Sajjanar, et al., 2024; Puvanendran, et al., 2023; Kelly, et al., 2020; Beemelmanns, et al., 2021). The GO enrichment analysis indicated that the set of DEGs was significantly enriched for specific biological functions and components, but interpreted with caution, because the function of many proteins and GO terms have been determined mainly through work in humans or mice. When GO terms represented biological processes, molecular functions, or cellular components seemingly irrelevant in fish (e.g. GO terms related to diseases), we excluded them from interpretation to maintain relevance for our study organism. The GO terms for which our set of DEGs was significantly enriched tended to have fewer genes associated with them than other fish studies (Puvanendran, et al., 2023), suggesting that the identified GO terms were more specific to particular biological processes, as the higher-level GO terms tend to encompass many genes. Upon entering anoxia, the predicted functions of the upregulated genes were primarily related to transcription regulation, including RNA polymerase II activity, DNA-templated transcription, and DNA binding, highlighting that the regulation of transcription could be important for survival under low oxygen conditions. Additional GO terms highlighted genes predicted to be associated with angiogenesis and vasculogenesis, which would be needed for new blood vessel formation to support an elevated glucose and lactate transport needed to sustain exclusively anaerobic metabolism. The upregulation of L-ascorbic acid binding, peptidyl-proline 4-dioxygenase activity, nucleus, and chromatin terms underscored a potential role of the antioxidant defense, collagen stabilization, and extensive transcriptional changes during anoxia. Overrepresented GO terms in the set of downregulated genes included fatty acid metabolism, which makes sense given the known necessary shift from oxygen-dependent ATP production to glycolysis as supported by metabolomics data (Dahl, Johansen, Nilsson, & Lefevre, 2021) and phospholipid homeostasis indicating modifications in membrane composition (Farhat, Turenne, Choi, & Weber, 2019). Conversely, the nucleolus GO term among downregulated genes suggests an association with reduced transcription and translation, potentially serving as a mechanism for energy conservation under anoxic conditions. Reduced transcriptional activity is corroborated by earlier findings of reduced RNA-to-protein ratio in crucian carp brain calculated from measured levels of injected labeled phenylalanine (Smith, Houlihan, Nilsson, & Brechin, 1996). The transition from anoxia to reoxygenation seems to involve extensive adjustments to the transcriptome, as indicated by overrepresentation of GO terms such as DNA-binding transcription factor activity, RNA polymerase II-specific, RNA polymerase II cis-regulatory region sequence-specific DNA binding, and double-stranded DNA binding. These terms might suggest a continuity in mRNA abundance changes from anoxia rather than a specific response to reoxygenation, as many factors were also present during the transition from normoxia to anoxia. Enrichment of signaling terms, such as C-C chemokine binding and C-C chemokine receptor activity, indicated that transmembrane signaling could be vital for the response to anoxia. The GO term cellular response to stress reflects various changes due to external stress. GO terms such as regulation of cell differentiation, establishment of epithelial cell polarity, cell maturation, neuronal cell population maintenance, and glial cell fate commitment suggest significant differentiation activities during reoxygenation. For instance, the GO term negative regulation of forebrain neuron differentiation could imply that neuron differentiation is stalled and points towards maintaining the current cells rather than generating new ones during initial reoxygenation.

By investigating the impact of DNA methylation on gene expression using WGBS, we identified 53 genes with DMRs within their gene borders across all comparisons: 14 genes for N to A, 28 for N to R, and 15 for A to R. Since the heatmap (Figure 7) demonstrated that all normoxia samples cluster together, it suggests that sex is not likely a confounding factor, as was also indicated by the PCA (Figure 3A). Consistently, the golden pompano fish (*Trachinotus blochii*) showed no significant difference between males and females in response to hypoxia and reoxygenation (Gu, et al., 2024). Compared to the hundreds of DMRs found in related studies, our study identified fewer DMRs, likely due to using the highly conservative, sample-independent program MethylScore (Zhang, et al., 2024). MethylScore avoids experimenter bias by not requiring predefined sample groups and ensures reliable results by identifying DMRs in at least 20% of samples. It uses a two-state hidden Markov model from WGBS data to avoid unreliable assumptions and arbitrary thresholds, reducing background noise (Hüther, et al., 2022). Hüther et al. (2022) notes that many DMR identification methods, like methylKit (Akalin, et al., 2012), create a multiple-testing burden by defining DMRs as clusters of adjacent methylated cytosines. In contrast, methods like metilene (Jühling, et al., 2015), HOME (Srivastava, Karpievitch, Eichten, Borevitz, & Lister, 2019), and dmrseq (Korthauer, Chakraborty, Benjamini, & Irizarry, 2018), which call DMRs only in predefined regions, may miss relevant loci. Furthermore, programs designed for mammalian DNA methylation data may not be suitable for non-mammalian sequence contexts, genomic distributions, and methylation patterns. MethylScore generalizes DMR calling across species with an unsupervised model trained on species-specific methylation (Hüther, et al., 2022). DNA methylation studies also use various methods for data preparation, which can affect comparisons. For example, Puvanendran et al. (2023) used reduced representation bisulfite sequencing (RRBS) (Puvanendran, et al., 2023), while we used genome-wide bisulfite sequencing. Differences also exist in library preparation kits (EZ DNA Methylation-Gold Kit (Wang, et al., 2022; Yue, et al., 2023), compared to our Pico Methyl-Seq™ Library Prep Kit) and methylation thresholds (e.g., 20% for Methylscore vs. 10-25% for Methylpy (Schultz, et al., 2015)) and criteria for contextualizing DMRs (e.g., proximity to genes: 2 kb (Puvanendran, et al., 2023), or 5 kb (Emmerson, Bechtold, Zabet, & Lawson, 2024)). These methodological differences can lead to the overestimation or underestimation of DMRs, complicating study comparisons.

Most DMRs in our study were located inside the transcribed region, with some before the transcriptional start and many overlapping it, aligning with existing literature (Katirtzoglou, et al., 2024; Ma, et al., 2024). The staircase plot revealed distinct clustering patterns of DMRs across comparisons, with some clustering more in the first 25% of gene length and others featuring longer DMRs, enhancing our understanding of DMR distribution. When evaluating methylation status, we observed a balance between hyper- and hypomethylation when entering anoxia, indicating the need to activate anoxia-specific genes and inactivate normoxia-specific ones. Upon reoxygenation, hypomethylation predominated, suggesting more genes need activation. It can be hypothesized that genes that were “turned off” in anoxia are most likely needed specifically for reoxygenation and are “turned back on” again when returning to normoxia. We demonstrated that for some specific genes hypermethylation leads to reduced mRNA abundance, and hypomethylation leads to gene activation around the transcriptional start site, consistent with existing literature (Kulis & Esteller, 2010). However, methylation levels upstream or downstream of the transcriptional start site showed positive and negative associations with gene expression, indicating a more complex pattern. Whereas our research examined adult crucian carp, most studies correlating DMRs and DEGs have focused on juvenile fish. Temperature effects were studied in Atlantic cod embryos (Puvanendran, et al., 2023), juvenile large yellow croakers were examined in hypoxic and acidic conditions (Yue, et al., 2023), adaptive plasticity was found in developing salmonids under heat and low oxygen (Kelly, et al., 2020), effects of atrazine exposure was demonstrated in zebrafish post-fertilization (Wang, et al., 2022), and hypoxia and oil exposure was explored in larval sheepshead minnows (Jones & Griffitt, 2022). All the mentioned studies found varying associations between DMRs and DEGs, suggesting that correlations between DMRs and DEGs may be more significant during early developmental stages. They concluded that DNA methylation changes are not the primary drivers of transcriptional patterns, as small correlations suggest a limited role in transcription (Jones & Griffitt, 2022; Wang, et al., 2022). Our focus was on acute stress response mechanisms rather than developmental aspects. However, 17 of the 53 DMR genes did show expression changes that were possibly regulated by DNA methylation, while other factors might be regulating mRNA abundance of other DEGs in anoxia and reoxygenation. The 17 DMR genes code for proteins with roles in transcriptional regulation, RNA splicing, protein trafficking, degradation, activity regulation, signal transduction, and cell integrity. Genes coding for proteins involved in DNA-level transcription regulation were identified, including ERF (ETS domain-containing transcription factor), a phosphorylation-modulated repressor of cell proliferation and cycle transition (Sgouras, et al., 1995), and RuvB-like 1, a DNA-binding protein essential for chromatin remodeling and gene transcription (Rottbauer, et al., 2002). Lysine-specific demethylase 1 plays dual roles in repressing and activating gene expression by demethylating histone H3 and reducing pro-inflammatory cytokine release to minimize tissue damage (Baby, Shinde, Kulkarni, & Sahu, 2023; Kim, Nam, & Baek, 2023). Telomerase Cajal body protein 1 is vital for telomere synthesis, highlighting the complexity of transcription regulation and cellular maintenance. The transcription of most genes that likely code for transcription-regulating proteins is downregulated during anoxia and is restored during reoxygenation (Figure 8 and Figure 9). ERF slightly increases from anoxia to reoxygenation, while Telomerase Cajal body protein 1 remains stable, suggesting prioritization of cell survival over genomic integrity. TIA1 and SRSF1 are key players in RNA splicing. TIA1 regulates alternative splicing and translation repression, forming stress granules under stress (Förch, et al., 2000; Kedersha, Gupta, Li, Miller, & Anderson, 1999), and is downregulated during anoxia. SRSF1, involved in spliceosome assembly and alternative splicing, shows increased expression during anoxia, indicating its potential role in fine-tuning stress response genes (Ning, Sandoval-Castellanos, Bhargava, Zhao, & Xu, 2023). The mRNA abundance of both proteins returns to normal levels upon reoxygenation. During anoxia, RNA splicing genes involved in transcription decreased, while upon reoxygenation, the transcription of the spliceosome and alternative splicing factors increased. Dynactin subunit 2 and PRA1 family protein 3, crucial for ER-Golgi vesicle trafficking, are downregulated during anoxia, indicating they may aid energy conservation through their role in translation (Tempes, Weslawski, Brzozowska, & Jaworski, 2020; Liu, Wu, & Chang, 2006). Upon reoxygenation, Dynactin subunit 2 partially restores, hinting at gradual functional reactivation, while PRA1 exceeds normoxia levels, implying a facilitative role for rapid recovery and reestablishing homeostasis. Proteins in the ubiquitin-proteasome system, such as E3 ubiquitin-protein ligase UBR2 (Kim, et al., 2021), F-box/LRR-repeat protein 20 (Jin, et al., 2004), and KCTD2 (Kim, Kim, Jin, Jin, & Kim, 2017), are downregulated during anoxia, indicating a reduced protein degradation rate and potential prioritization of protein stability for essential functions. UBR2 recovers upon reoxygenation, while F-box/LRR-repeat protein 20 and KCTD2 show partial recovery. SPAT2, involved in protein de-ubiquitination and neuroprotection, is also downregulated during anoxia and resumes expression upon reoxygenation. This suggests it may serve as an energy-conservation mechanism during low oxygen conditions and potentially aid recovery after anoxic damage (Moro, et al., 2007; Ren, et al., 2021). The Prolyl endopeptidase regulates protein activity by cleaving peptides at the C-terminal side of proline residues and is downregulated during anoxia and reoxygenation, which may reflect a preventive mechanism to avoid neuropeptide breakdown. However, its exact physiological role remains unclear (Wilk, 1983; Vanhoof, et al., 1994). During anoxia, GRB2 (Growth Factor Receptor-Bound Protein 2) and LRP12 (Low-density Lipoprotein Receptor-related Protein 12), critical in growth factor receptor pathways and endocytosis, were downregulated, implying that cells may minimize non-essential signaling to conserve resources (Malagrinò, Puglisi, Pagano, Travaglini-Allocatelli, & Toto, 2024; Battle, Maher, & McCormick, 2003). As reoxygenation occurs, the mRNA abundance of these proteins gradually recovered, suggesting the facilitation of normal cellular functions and aiding overall recovery. Septin-9 and Calmin are crucial for cell structure and function. Septin-9, a cytoskeletal GTPase involved in cytokinesis, cell polarization, vesicle trafficking, and apoptosis, was downregulated during anoxia, hinting at a halt of cell division, which would aid in conserving resources. Sun et al. (2020) described that overexpression of Septin-9 inhibits the degradation of HIF-1-alpha, leading to increased VEGF expression and enhanced angiogenesis (Sun, Zheng, Li, & Zhang, 2020). Upon reoxygenation, Septin-9 mRNA expression exceeded normoxic levels, indicating that stressed cells may overexpress Septin-9 to restore tissue oxygen supply rapidly. Calmin, which connects the ER to actin fibers, showed increased mRNA expression during both anoxia and reoxygenation, suggesting that maintaining cell integrity under stress may be necessary (Ishisaki, Takaishi, Furuta, & Huh, 2001; Merta, Isogai, Paul, Danuser, & Henne, 2024).

Taken together our study suggests that anoxia induces changes in DNA methylation patterns of specific genes and, therefore, may play a role in regulating the response to anoxia in crucian carp brains. The 17 genes exhibiting both differential methylation and altered mRNA abundance as determined by independent analyses appeared to play roles in transcriptional regulation, RNA splicing, protein trafficking, degradation, activity regulation, signal transduction, and maintaining cell integrity. Through transcriptome analyses, we identified hundreds of differentially expressed genes with putative roles in gene regulation, vascularization, and immune response. Further exploration of higher-level epigenomic mechanisms - including chromatin accessibility through ATAC sequencing and histone modifications such as acetylation, phosphorylation, and ubiquitination - could enhance our comprehension of how crucian carp cope with seasonal and prolonged periods of oxygen deprivation. However, these modifications may be more stable than DNA methylation patterns, so whether they play a role in the mechanisms of anoxia tolerance in a seasonal context remains uncertain. This work advances our understanding of epigenomic changes under extreme conditions, offering insights into oxygen-stress responses and low-oxygen adaptation.

## Author contribution

**Magdalena Winklhofer**: Software, Formal analysis, Investigation, Data curation, Visualization, Writing-Original Draft, Writing-Review and Editing; **Øivind Andersen**: Conceptualization (supporting), Supervision (supporting), Writing-Review and Editing (supporting); **Sjannie Lefevre**: Conceptualization (lead), Methodology, Resources, Writing-Review and Editing (lead), Supervision (lead), Project administration, Funding acquisition.

## Data availability

All raw sequencing data are deposited in the NCBI Sequence Read Archive (SRA) under BioProject ID PRJNA1163668 (http://www.ncbi.nlm.nih.gov/bioproject/1163668). The raw genome sequences are available in SRA (BioProject: PRJNA1119394), and the assembly and annotation data were obtained from DataverseNO (https://doi.org/10.18710/GXMSUH). Supplementary Material for the present study has been deposited in DataverseNO: https://doi.org/10.18710/GSHJEB, and scripts are available in the GitHub repository WholeGenomeBisulphiteSequencing (https://github.com/MagdalenaWinklhofer/WholeGenomeBisulphiteSequencing.git).

## Acknowledgments

The authors thank the staff at the Norwegian Sequencing Center for preparing the RNA library preparation, and DNA and RNA sequencing; Sigma2 – the National Infrastructure of High-Performance Computing and Data Storage in Norway for enabling data analysis using the high-performance computing cluster Saga; and the staff at the InVivo facility at the Department of Biosciences for supporting crucian carp husbandry. The authors also thank the Lefevre-Nilsson research group member Lucie Gerber and former members Tellef Helle-Valle and Helge-Andre Dahl, for their assistance with fish collection, anoxia exposure, and sampling. Special thanks are also given to Göran E. Nilsson for insightful discussions.

## References

Akalin, A., Kormaksson, M., Li, S., Garrett-Bakelman, F. E., Figueroa, M. E., Melnick, A., & Mason, C. E. (2012, October). methylKit: a comprehensive R package for the analysis of genome-wide DNA methylation profiles. Genome Biology, 13. doi:10.1186/gb-2012-13-10-r87

Baby, S., Shinde, S. D., Kulkarni, N., & Sahu, B. (2023, October). Lysine-Specific Demethylase 1 (LSD1) Inhibitors: Peptides as an Emerging Class of Therapeutics. ACS Chemical Biology, 18, 2144– 2155. doi:10.1021/acschembio.3c00386

Battle, M. A., Maher, V. M., & McCormick, J. J. (2003, May). ST7 Is a Novel Low-Density Lipoprotein Receptor-Related Protein (LRP) with a Cytoplasmic Tail that Interacts with Proteins Related to Signal Transduction Pathways. Biochemistry, 42, 7270–7282. doi:10.1021/bi034081y

Beemelmanns, A., Ribas, L., Anastasiadi, D., Moraleda-Prados, J., Zanuzzo, F. S., Rise, M. L., & Gamperl, A. K. (2021, January). DNA Methylation Dynamics in Atlantic Salmon (Salmo salar) Challenged With High Temperature and Moderate Hypoxia. Frontiers in Marine Science, 7. doi:10.3389/fmars.2020.604878

Boutilier, R. G. (2001, September). Mechanisms of cell survival in hypoxia and hypothermia. Journal of Experimental Biology, 204, 3171–3181. doi:10.1242/jeb.204.18.3171

Dahl, H.-A., Johansen, A., Nilsson, G. E., & Lefevre, S. (2021, July 1). The Metabolomic Response of Crucian Carp (Carassius carassius) to Anoxia and Reoxygenation Differs between Tissues and Hints at Uncharacterized Survival Strategies. Metabolites, 11, 435. doi:10.3390/metabo11070435

Dupont, C., Armant, D., & Brenner, C. (2009, August). Epigenetics: Definition, Mechanisms and Clinical Perspective. Seminars in Reproductive Medicine, 27, 351–357. doi:10.1055/s-0029-1237423

Emmerson, R. A., Bechtold, U., Zabet, N. R., & Lawson, T. (2024). DNA methylation contributes to plant acclimation to naturally fluctuating light. bioRxiv preprint doi: 10.1101/2024.06.07.597890; this version posted June 9, 2024. The copyright holder for this preprint (which was not certified by peer review) is the author/funder, who has granted bioRxiv a license to display the preprint in perpetuity. It is made.

Erecińska, M., & Silver, I. A. (1994, May). Ions and energy in mammalian brain. Progress in Neurobiology, 43, 37–71. doi:10.1016/0301-0082(94)90015-9

Esteller, M. (2008, March). Epigenetics in Cancer. New England Journal of Medicine, 358, 1148–1159. doi:10.1056/nejmra072067

Farhat, E., Turenne, E. D., Choi, K., & Weber, J.-M. (2019, November). Hypoxia-induced remodelling of goldfish membranes. Comparative Biochemistry and Physiology Part B: Biochemistry and Molecular Biology, 237, 110326. doi:10.1016/j.cbpb.2019.110326

Feil, R., Charlton1, J., Bird1, A., Walter, J., & Reik, W. (1994). Methylation analysis on individual chromosomes: improved protocol for bisulphite genomic sequencing.

Feng, S., Jacobsen, S. E., & Reik, W. (2010, October). Epigenetic Reprogramming in Plant and Animal Development. Science, 330, 622–627. doi:10.1126/science.1190614

Förch, P., Puig, O., Kedersha, N., Martínez, C., Granneman, S., Séraphin, B., . . . Valcárcel, J. (2000, November). The Apoptosis-Promoting Factor TIA-1 Is a Regulator of Alternative Pre-mRNA Splicing. Molecular Cell, 6, 1089–1098. doi:10.1016/s1097-2765(00)00107-6

Gong, T., Borgard, H., Zhang, Z., Chen, S., Gao, Z., & Deng, Y. (2022, January). Analysis and Performance Assessment of the Whole Genome Bisulfite Sequencing Data Workflow: Currently Available Tools and a Practical Guide to Advance DNA Methylation Studies. Small Methods, 6. doi:10.1002/smtd.202101251

Gu, Y., Jin, C. X., Tong, Z. H., Jiang, T., Yao, F. C., Zhang, Y., . . . Luo, J. (2024, July). Expression of genes related to gonadal development and construction of gonadal DNA methylation maps of Trachinotus blochii under hypoxia. Science of The Total Environment, 935, 173172. doi:10.1016/j.scitotenv.2024.173172

Haverinen, J., Badr, A., Eskelinen, M., & Vornanen, M. (2024, January). Three steps down: Metabolic depression in winter-acclimatized crucian carp (Carassius carassius L.). Comparative Biochemistry and Physiology Part A: Molecular & Integrative Physiology, 287, 111537. doi:10.1016/j.cbpa.2023.111537

Holliday, R. (1987, October). The Inheritance of Epigenetic Defects. Science, 238, 163–170. doi:10.1126/science.3310230

Holopainen, A. I., Tonn, W. M., & Paszkowski, C. A. (1997). Tales of two fish: the dichotomous biology of crucian carp (Carassius carassius (L.)) in northern Europe.

Holopainen, I. J., Hyvärinen, H., & Piironen, J. (1986, January). Anaerobic wintering of crucian carp (Carassius carassius L.)—II. Metabolic products. Comparative Biochemistry and Physiology Part A: Physiology, 83, 239–242. doi:10.1016/0300-9629(86)90568-2

Hüther, P., Hagmann, J., Nunn, A., Kakoulidou, I., Pisupati, R., Langenberger, D., . . . Becker, C. (2022). MethylScore, a pipeline for accurate and context-aware identification of differentially methylated regions from population-scale plant whole-genome bisulfite sequencing data. Quantitative Plant Biology, 3. doi:10.1017/qpb.2022.14

Ishisaki, Z., Takaishi, M., Furuta, I., & Huh, N.-h. (2001, June). Calmin, a Protein with Calponin Homology and Transmembrane Domains Expressed in Maturing Spermatogenic Cells. Genomics, 74, 172–179. doi:10.1006/geno.2001.6544

Jin, J., Cardozo, T., Lovering, R. C., Elledge, S. J., Pagano, M., & Harper, J. W. (2004, November). Systematic analysis and nomenclature of mammalian F-box proteins. Genes & Development, 18, 2573–2580. doi:10.1101/gad.1255304

Johansson, D., Nilsson, G. E., & Törnblom, E. (1995, March). Effects of Anoxia on Energy Metabolism in Crucian Carp Brain Slices Studied With Microcalorimetry. Journal of Experimental Biology, 198, 853–859. doi:10.1242/jeb.198.3.853

Johnston, I., & Bernard, L. (1983). Utilization of the ethanol pathway in carp following exposure to anoxia.

Jones, E. R., & Griffitt, R. J. (2022, October). Oil and hypoxia alter DNA methylation and transcription of genes related to neurological function in larval Cyprinodon variegatus. Aquatic Toxicology, 251, 106267. doi:10.1016/j.aquatox.2022.106267

Jones, P. A. (2012, May). Functions of DNA methylation: islands, start sites, gene bodies and beyond. Nature Reviews Genetics, 13, 484–492. doi:10.1038/nrg3230

Jühling, F., Kretzmer, H., Bernhart, S. H., Otto, C., Stadler, P. F., & Hoffmann, S. (2015, December). metilene: fast and sensitive calling of differentially methylated regions from bisulfite sequencing data. Genome Research, 26, 256–262. doi:10.1101/gr.196394.115

Jurcau, A., & Simion, A. (2021, December). Neuroinflammation in Cerebral Ischemia and Ischemia/Reperfusion Injuries: From Pathophysiology to Therapeutic Strategies. International Journal of Molecular Sciences, 23, 14. doi:10.3390/ijms23010014

Katirtzoglou, A., Hansen, S. B., Sveier, H., Martin, M. D., Brealey, J. C., & Limborg, M. T. (2024, August). Genomic context determines the effect of DNA methylation on gene expression in the gut epithelium of Atlantic salmon (Salmo salar). Epigenetics, 19. doi:10.1080/15592294.2024.2392049

Kedersha, N. L., Gupta, M., Li, W., Miller, I., & Anderson, P. (1999, December). RNA-Binding Proteins Tia-1 and Tiar Link the Phosphorylation of Eif-2α to the Assembly of Mammalian Stress Granules. The Journal of Cell Biology, 147, 1431–1442. doi:10.1083/jcb.147.7.1431

Kelly, T., Johnsen, H., Burgerhout, E., Tveiten, H., Thesslund, T., Andersen, Ø., & Robinson, N. (2020, September 3). Low Oxygen Stress During Early Development Influences Regulation of Hypoxia-Response Genes in Farmed Atlantic Salmon (Salmo salar). G3 Genes Genomes Genetics, 10, 3179–3188. doi:10.1534/g3.120.401459

Kim, D., Nam, H. J., & Baek, S. H. (2023, December). Post-translational modifications of lysine-specific demethylase 1. Biochimica et Biophysica Acta (BBA) - Gene Regulatory Mechanisms, 1866, 194968. doi:10.1016/j.bbagrm.2023.194968

Kim, E.-J., Kim, S.-H., Jin, X., Jin, X., & Kim, H. (2017, January). KCTD2, an adaptor of Cullin3 E3 ubiquitin ligase, suppresses gliomagenesis by destabilizing c-Myc. Cell Death & Differentiation, 24, 649–659. doi:10.1038/cdd.2016.151

Kim, J. G., Shin, H.-C., Seo, T., Nawale, L., Han, G., Kim, B. Y., . . . Cha-Molstad, H. (2021, August). Signaling Pathways Regulated by UBR Box-Containing E3 Ligases. International Journal of Molecular Sciences, 22, 8323. doi:10.3390/ijms22158323

Korthauer, K., Chakraborty, S., Benjamini, Y., & Irizarry, R. A. (2018, February). Detection and accurate false discovery rate control of differentially methylated regions from whole genome bisulfite sequencing. Biostatistics, 20, 367–383. doi:10.1093/biostatistics/kxy007

Kristián, T., & Siesjö, B. K. (1996, July). Calcium-related damage in ischemia. Life Sciences, 59, 357–367. doi:10.1016/0024-3205(96)00314-1

Kulis, M., & Esteller, M. (2010). DNA Methylation and Cancer. In Advances in Genetics (pp. 27–56). Elsevier. doi:10.1016/b978-0-12-380866-0.60002-2

Lefevre, S., & Nilsson, G. E. (2023, April). Two decades of research on anoxia tolerance – mitochondria, -omics and physiological diversity. Journal of Experimental Biology, 226. doi:10.1242/jeb.245584

Lefevre, S., & Nilsson, G. E. (2024). Case study: The anoxia-tolerant crucian carp. In Encyclopedia of Fish Physiology (pp. 148–158). Elsevier. doi:10.1016/b978-0-323-90801-6.00105-1

Lefevre, S., Stecyk, J. A., Torp, M.-K., Løvold, L. Y., Sørensen, C., Johansen, I. B., . . . Nilsson, G. E. (2017, November). Re-oxygenation after anoxia induces brain cell death and memory loss in the anoxia-tolerant crucian carp. Journal of Experimental Biology, 220, 3883–3895. doi:10.1242/jeb.165118

Li, Y. (2021, March). Modern epigenetics methods in biological research. Methods, 187, 104–113. doi:10.1016/j.ymeth.2020.06.022

Liu, H.-P., Wu, C.-C., & Chang, Y.-S. (2006, August). PRA1 promotes the intracellular trafficking and NF-κB signaling of EBV latent membrane protein 1. The EMBO Journal, 25, 4120–4130. doi:10.1038/sj.emboj.7601282

Lucena-Aguilar, G., Sánchez-López, A. M., Barberán-Aceituno, C., Carrillo-Ávila, J. A., López-Guerrero, J. A., & Aguilar-Quesada, R. (2016, August). DNA Source Selection for Downstream Applications Based on DNA Quality Indicators Analysis. Biopreservation and Biobanking, 14, 264–270. doi:10.1089/bio.2015.0064

Lutz, P. L., & Nilsson, G. E. (1997, January). Contrasting Strategies for Anoxic Brain Survival – Glycolysis Up or Down. Journal of Experimental Biology, 200, 411–419. doi:10.1242/jeb.200.2.411

Ma, J., Shi, K., Zhang, W., Han, S., Wu, Z., Wang, M., . . . Sha, Z. (2024, September). The survival, gene expression, and DNA methylation of Paralichthys olivaceus impacted by the decay of green tide and bacterial infection in both laboratory and field simulation experiments. Science of The Total Environment, 942, 173427. doi:10.1016/j.scitotenv.2024.173427

Malagrinò, F., Puglisi, E., Pagano, L., Travaglini-Allocatelli, C., & Toto, A. (2024, September). GRB2: A dynamic adaptor protein orchestrating cellular signaling in health and disease. Biochemistry and Biophysics Reports, 39, 101803. doi:10.1016/j.bbrep.2024.101803

Merta, H., Isogai, T., Paul, B., Danuser, G., & Henne, W. M. (2024, January). Spatial proteomics of ER tubules reveals CLMN, an ER-actin tether at focal adhesions that promotes cell migration. doi:10.1101/2024.01.24.577043

Moro, E., Maran, C., Slongo, M. L., Argenton, F., Toppo, S., & Onisto, M. (2007). Zebrafish spata2 is expressed at early developmental stages. The International Journal of Developmental Biology, 51, 241–246. doi:10.1387/ijdb.062220em

Nilsson, G. E. (1990, May). Long-term anoxia in crucian carp: changes in the levels of amino acid and monoamine neurotransmitters in the brain, catecholamines in chromaffin tissue, and liver glycogen. Journal of Experimental Biology, 150, 295–320. doi:10.1242/jeb.150.1.295

Ning, K., Sandoval-Castellanos, A., Bhargava, A., Zhao, M., & Xu, J. (2023). Serine and arginine rich splicing factor 1: a potential target for neuroprotection and other diseases. Neural Regeneration Research, 18, 1411. doi:10.4103/1673-5374.360243

Puvanendran, V., Burgerhout, E., Andersen, Ø., Kent, M., Hansen, Ø., & Tengs, T. (2023, July). Intergenerational effects of early life-stage temperature modulation on gene expression and {DNA} methylation in Atlantic cod ($\less$i$\greater$Gadus morhua$\less$/i$\greater$). Epigenetics, 18. doi:10.1080/15592294.2023.2237759

Qin, C., Yang, S., Chu, Y.-H., Zhang, H., Pang, X.-W., Chen, L., . . . Wang, W. (2022, July). Signaling pathways involved in ischemic stroke: molecular mechanisms and therapeutic interventions. Signal Transduction and Targeted Therapy, 7. doi:10.1038/s41392-022-01064-1

Ren, Y., Jiang, J., Jiang, W., Zhou, X., Lu, W., Wang, J., & Luo, Y. (2021, June). Spata2 Knockdown Exacerbates Brain Inflammation via NF-κB/P38MAPK Signaling and NLRP3 Inflammasome Activation in Cerebral Ischemia/Reperfusion Rats. Neurochemical Research, 46, 2262–2275. doi:10.1007/s11064-021-03360-8

Riggs, C. L., Summers, A., Warren, D. E., Nilsson, G. E., Lefevre, S., Dowd, W. W., . . . Podrabsky, J. E. (2018, July). Small Non-coding RNA Expression and Vertebrate Anoxia Tolerance. Frontiers in Genetics, 9. doi:10.3389/fgene.2018.00230

Robinson, M. D., & Oshlack, A. (2010). A scaling normalization method for differential expression analysis of RNA-seq data. Genome Biology, 11, R25. doi:10.1186/gb-2010-11-3-r25

Rottbauer, W., Saurin, A. J., Lickert, H., Shen, X., Burns, C. G., Wo, Z. G., . . . Fishman, M. (2002, November). Reptin and Pontin Antagonistically Regulate Heart Growth in Zebrafish Embryos. Cell, 111, 661–672. doi:10.1016/s0092-8674(02)01112-1

Ruhr, I., Bierstedt, J., Rhen, T., Das, D., Singh, S. K., Miller, S., . . . Galli, G. L. (2021, September). Developmental programming of DNA methylation and gene expression patterns is associated with extreme cardiovascular tolerance to anoxia in the common snapping turtle. Epigenetics & Chromatin, 14. doi:10.1186/s13072-021-00414-7

Sajjanar, B., Aalam, M. T., Khan, O., Dhara, S. K., Ghosh, J., Gandham, R. K., . . . Mishra, B. P. (2024, August). Genome-wide DNA methylation profiles regulate distinct heat stress response in zebu (Bos indicus) and crossbred (Bos indicus × Bos taurus) cattle. Cell Stress and Chaperones, 29, 603–614. doi:10.1016/j.cstres.2024.06.005

Saxonov, S., Berg, P., & Brutlag, D. L. (2006, January). A genome-wide analysis of CpG dinucleotides in the human genome distinguishes two distinct classes of promoters. Proceedings of the National Academy of Sciences, 103, 1412–1417. doi:10.1073/pnas.0510310103

Schultz, M. D., He, Y., Whitaker, J. W., Hariharan, M., Mukamel, E. A., Leung, D., . . . Ecker, J. R. (2015, June). Human body epigenome maps reveal noncanonical DNA methylation variation. Nature, 523, 212–216. doi:10.1038/nature14465

Schwartz, S., Meshorer, E., & Ast, G. (2009, August). Chromatin organization marks exon-intron structure. Nature Structural & Molecular Biology, 16, 990–995. doi:10.1038/nsmb.1659

Sgouras, D. N., Athanasiou, M. A., Beal, G. J., Fisher, R. J., Blair, D. G., & Mavrothalassitis, G. J. (1995, October). ERF: an ETS domain protein with strong transcriptional repressor activity, can suppress ets-associated tumorigenesis and is regulated by phosphorylation during cell cycle and mitogenic stimulation. The EMBO Journal, 14, 4781–4793. doi:10.1002/j.1460-2075.1995.tb00160.x

Smith, R. W., Houlihan, D. F., Nilsson, G. E., & Brechin, J. G. (1996, October). Tissue-specific changes in protein synthesis rates in vivo during anoxia in crucian carp. American Journal of Physiology-Regulatory, Integrative and Comparative Physiology, 271, R897–R904. doi:10.1152/ajpregu.1996.271.4.r897

Smith, Z. D., & Meissner, A. (2013, February). DNA methylation: roles in mammalian development. Nature Reviews Genetics, 14, 204–220. doi:10.1038/nrg3354

Srivastava, A., Karpievitch, Y. V., Eichten, S. R., Borevitz, J. O., & Lister, R. (2019, May). HOME: a histogram based machine learning approach for effective identification of differentially methylated regions. BMC Bioinformatics, 20. doi:10.1186/s12859-019-2845-y

Struhl, K. (2024, December). The distinction between epigenetics and epigenomics. Trends in Genetics, 40, 995–997. doi:10.1016/j.tig.2024.10.002

Sun, J., Zheng, M.-Y., Li, Y.-W., & Zhang, S.-W. (2020, June). Structure and function of Septin 9 and its role in human malignant tumors. World Journal of Gastrointestinal Oncology, 12, 619–631. doi:10.4251/wjgo.v12.i6.619

Sun, L., Luo, H., Bu, D., Zhao, G., Yu, K., Zhang, C., . . . Zhao, Y. (2013, July). Utilizing sequence intrinsic composition to classify protein-coding and long non-coding transcripts. Nucleic Acids Research, 41, e166–e166. doi:10.1093/nar/gkt646

Supek, F., Bošnjak, M., Škunca, N., & Šmuc, T. (2011, July). REVIGO Summarizes and Visualizes Long Lists of Gene Ontology Terms. (C. Gibas, Ed.) PLoS ONE, 6, e21800. doi:10.1371/journal.pone.0021800

Tempes, A., Weslawski, J., Brzozowska, A., & Jaworski, J. (2020, April). Role of dynein–dynactin complex, kinesins, motor adaptors, and their phosphorylation in dendritogenesis. Journal of Neurochemistry, 155, 10–28. doi:10.1111/jnc.15010

Valencia-Pesqueira, L. M., Hoff, S. N., Tørresen, O. K., Jentoft, S., & Lefevre, S. (2025, March). Chromosome-level de novo genome assembly of wild, anoxia-tolerant crucian carp, Carassius carassius. Scientific Data, 12. doi:10.1038/s41597-025-04813-3

Valencia-Pesqueira, L. M., Hoff, S. N., Tørresen, O. K., Jentoft, S., & Lefevre, S. (2025). Replication Data for: Chromosome-level de novo genome assembly of wild, anoxia-tolerant crucian carp, Carassius carassius. Replication Data for: Chromosome-level de novo genome assembly of wild, anoxia-tolerant crucian carp, Carassius carassius. DataverseNO. doi:10.18710/GXMSUH

Vanhoof, G., Goossens, F., Hendriks, L., De Meester, I., Hendriks, D., Vriend, G., . . . Scharpé, S. (1994, November). Cloning and sequence analysis of the gene encoding human lymphocyte prolyl endopeptidase. Gene, 149, 363–366. doi:10.1016/0378-1119(94)90177-5

Vornanen, M., Stecyk, J. A., & Nilsson, G. E. (2009). Chapter 9 The Anoxia-Tolerant Crucian Carp (Carassius Carassius L.). In Fish Physiology (pp. 397–441). Elsevier. doi:10.1016/s1546-5098(08)00009-5

Waechter, D. E., & Baserga, R. (1982, February). Effect of methylation on expression of microinjected genes. Proceedings of the National Academy of Sciences, 79, 1106–1110. doi:10.1073/pnas.79.4.1106

Wang, S., Bryan, C., Xie, J., Zhao, H., Lin, L. F., Tai, J. A., . . . Yuan, C. (2022, July). Atrazine exposure in zebrafish induces aberrant genome-wide methylation. Neurotoxicology and Teratology, 92, 107091. doi:10.1016/j.ntt.2022.107091

Wijenayake, S., & Storey, K. B. (2016, February). The role of DNA methylation during anoxia tolerance in a freshwater turtle (Trachemys scripta elegans). Journal of Comparative Physiology B, 186, 333–342. doi:10.1007/s00360-016-0960-x

Wilk, S. (1983, November). Prolyl endopeptidase. Life Sciences, 33, 2149–2157. doi:10.1016/0024-3205(83)90285-0

Wu, M.-Y., Yiang, G.-T., Liao, W.-T., Tsai, A.-Y., Cheng, Y.-L., Cheng, P.-W., . . . Li, C.-J. (2018). Current Mechanistic Concepts in Ischemia and Reperfusion Injury. Cellular Physiology and Biochemistry, 46, 1650–1667. doi:10.1159/000489241

Xie, J., Kittur, F., Li, P., & Hung, C.-Y. (2022). Rethinking the necessity of low glucose intervention for cerebral ischemia/reperfusion injury. Neural Regeneration Research, 17, 1397. doi:10.4103/1673-5374.330592

Yue, Y., Wang, Y., Zhang, B., Zeng, J., Wang, Q., Wang, C., & Peng, S. (2023, July). Whole-Genome Methylation Sequencing of Large Yellow Croaker (Larimichthys crocea) Liver Under Hypoxia and Acidification Stress. Marine Biotechnology. doi:10.1007/s10126-023-10226-3

Zhang, W., Li, X., Yu, U., Huang, X., Wang, H., Lu, Y., . . . Zhang, J. (2024, August). Genome-wide methylation and gene-expression analyses in thalassemia. Aging. doi:10.18632/aging.206037

Zhao, Y., Zhang, X., Chen, X., & Wei, Y. (2021, December). Neuronal injuries in cerebral infarction and ischemic stroke: From mechanisms to treatment (Review). International Journal of Molecular Medicine, 49. doi:10.3892/ijmm.2021.5070

Ziller, M. J., Hansen, K. D., Meissner, A., & Aryee, M. J. (2014, November). Coverage recommendations for methylation analysis by whole-genome bisulfite sequencing. Nature Methods, 12, 230–232. doi:10.1038/nmeth.3152

